# Loss of KMT2D accelerates hypertrophic chondrocyte differentiation and senescence by increasing mitochondrial ROS production

**DOI:** 10.64898/2026.03.18.712470

**Authors:** Sara Tholl Halldorsdottir, Stefán Pétursson, Jill A. Fahrner, Agnes Ulfig, Hans Tomas Bjornsson

## Abstract

Longitudinal bone growth occurs through endochondral ossification, which is accompanied by the differentiation of chondrocytes in the growth plate. Disruption in chondrocyte maturation can lead to skeletal growth abnormalities, such as those observed in Kabuki syndrome type 1 (KS1), a genetic disorder caused by heterozygous pathogenic variants in the *KMT2D* gene. KS1 patients exhibit postnatal growth deficiency, craniofacial hypoplasia, and skeletal deformities, yet the mechanisms underlying these phenotypic manifestations remain poorly understood. Our study investigated the effects of KMT2D deficiency on chondrocyte maturation and identified premature chondrocyte hypertrophy as a key driver of skeletal abnormalities in KS1. We previously observed reduced femur and tibia length in a KS1 mouse model, along with altered growth plate architecture, particularly affecting the heights of the proliferative and hypertrophic zones. Here, we show that KMT2D-deficient chondrocytes exhibit accelerated differentiation and early senescence upon exposure to supraphysiological oxygen levels (20% O_2_). These pathological changes were linked to increased mitochondrial reactive oxygen species (ROS) production likely caused by deficiencies in electron transport chain function, leading to oxidative stress and premature hypertrophy. Pharmacological ROS neutralization or hypoxic conditions mitigated these effects, restoring normal chondrocyte differentiation and preventing premature ossification. These findings demonstrate that KMT2D loss induces oxidative stress-driven chondrocyte hypertrophy, disrupting the balance of cartilage growth and ossification. Our study provides crucial mechanistic insights into KS1-associated skeletal abnormalities and suggests mitochondrial ROS regulation as a potential therapeutic avenue.

## Introduction

Longitudinal growth of long bones occurs through endochondral ossification, a process which is tightly regulated by the activity of the cartilage growth plate (reviewed in (Blumer, 2021)). This structure comprises three zones, the resting, proliferative, and hypertrophic zone, each containing chondrocytes at a different stage of differentiation. Chondrocytes in the resting zone, located closest to the epiphysis, serve as progenitor cells that generate new clones of rapidly proliferating chondrocytes, which form columns aligned with the bone’s long axis and secrete cartilage-specific extracellular matrix proteins, such as type II collagen and aggrecan (Long & Ornitz, 2013). As these cells divide, the daughter cells remain aligned to maintain this columnar structure. Chondrocytes further from the epiphysis stop proliferating, enlarge, and form the hypertrophic zone. These hypertrophic chondrocytes alter the cartilage matrix by secreting type X collagen (COLX) and higher levels of fibronectin to promote its calcification or mineralization. Terminally differentiated, late hypertrophic chondrocytes, farthest from the epiphysis, produce matrix metalloproteinases (MMPs) and release vascular endothelial growth factor (VEGF) before undergoing apoptosis, accelerating the degradation of the cartilage matrix and encouraging blood vessels and differentiating osteoblasts from the metaphysis to invade the hypertrophic zone. These bone-forming cells eventually remodel the cartilage into bone tissue, resulting in the formation of new bone underneath the growth plate, the so-called *primary spongiosa*, and consequently bone elongation (Araldi & Schipani, 2010; Blumer, 2021). While chondrocyte hypertrophy is mandatory for vascular invasion, osteoblast differentiation, and endochondral ossification (Blumer, 2021), inappropriate, i.e. excessive, premature, or blocked hypertrophy can cause skeletal growth abnormalities such as dwarfism (Arnold et al., 2007; Vega et al., 2004). Hence, control of chondrocyte hypertrophy is critical for normal longitudinal bone development.

The histone methyltransferase KMT2D is a core component of the COMPASS complex that is responsible for histone 3 lysine 4 (H3K4) mono-, di-, and tri-methylation, epigenetic modifications associated with active transcription (Froimchuk et al., 2017). Heterozygous pathogenic variants in the *KMT2D* gene are known to cause Kabuki syndrome type 1 (KS1), a rare genetic, multisystem disorder with a spectrum of clinical features including intellectual disability, craniofacial hypoplasia, skeletal deformities, and reduced postnatal growth (Adam et al., 2019). Given that most of these phenotypes are caused by deficiencies in the formation of cartilage and bones, KMT2D is thought to be involved in the regulation of endochondral ossification. However, the mechanisms by which KMT2D coordinates chondrocyte proliferation and differentiation during endochondral bone development remain elusive.

Intriguingly, our previous studies using a mouse model to evaluate growth abnormalities in KS1 revealed that in addition to having shortened femurs and tibias, the mice exhibited clear differences in the height of the proliferative and hypertrophic zone of their epiphyseal growth plate (Fahrner et al., 2019), indicating that the process of chondrocyte maturation might be altered in KS1. Follow-up *in vitro* studies using the chondrogenic cell line ATDC5 confirmed aberrant chondrocyte differentiation upon the loss of KMT2D, which manifested in a premature transition of the cells from a proliferative to a hypertrophic state (Fahrner et al., 2019). Such pathological changes in the growth plate and chondrogenesis could explain the observed disruption of the normal endochondral ossification process in KS1.

Although the epiphyseal growth plate undergoes neovascularization during endochondral bone formation, it remains an overall avascular tissue with a gradient of oxygenation from the proliferative to the hypertrophic zone that is correlated with the expression of the transcription factor hypoxia inducible factor 1 alpha (*HIF1A*) (Schipani et al., 2001). Proliferating chondrocytes reside in the most hypoxic microenvironment and critically depend on HIF1A for their survival, which stimulates anaerobic glycolysis while suppressing mitochondrial respiration and the production of mitochondrial reactive oxygen species (ROS) (Yao et al., 2019). Owing to the increased oxygen availability near the primary spongiosa and their overall higher energy demands, cells in the hypertrophic zone undergo a metabolic shift towards increasing mitochondrial respiration that leads to higher ROS production (Morita et al., 2007), facilitating their removal through apoptosis and replacement by osteoblasts during endochondral ossification. Yet, excessive ROS levels can also lead to oxidative stress and induce early cellular senescence which can adversely affect the balance between chondrocyte proliferation, maturation, and terminal differentiation, resulting in impaired longitudinal bone growth (Morita et al., 2007).

Interestingly, the growth deficiency in KS1 becomes evident not significantly earlier than in the postnatal period (Schott et al., 2016), with the intrauterine embryonic and fetal growth occurring largely normally until just before birth. This observation made us wonder whether an increased oxygen supply to the late fetal/postnatal growth plate and the resulting increase in mitochondrial ROS production could be causative for the premature chondrocyte hypertrophy in KS1.

In this study we aimed to gain mechanistic insights into how the loss of KMT2D drives precocious differentiation of chondrocytes, causing early onset chondrocyte hypertrophy and the dysregulation of the normal endochondral ossification process. We found that exposure of KMT2D-deficient chondrocytes to supraphysiological oxygen levels, i.e. 20% O_2_, resulted in accelerated differentiation and early senescence of hypertrophic chondrocytes which was accompanied by premature ossification. The increased sensitivity to oxygen likely resulted from deficiencies in the mitochondrial electron transport chain, leading to increased mitochondrial ROS production that exceeded a critical threshold in the intracellular ROS level, shifting the fate of the hypertrophic chondrocytes towards early senescence. Both the neutralization of mitochondrial ROS and the reduction of the oxygen level slowed down the pace of differentiation of KMT2D-deficient chondrocytes and inhibited premature hypertrophy. Our findings reveal a novel link between KMT2D deficiency, oxygen sensitivity, and chondrocyte maturation, suggesting that oxidative stress-induced hypertrophy contributes to disrupted endochondral ossification in KS1. These insights enhance our understanding of skeletal dysplasia pathogenesis and open up a novel approach of potential therapeutic strategies targeting mitochondrial ROS regulation.

## Results

### KMT2D-deficient chondrogenic cells are prone to differentiate and exit the cell cycle prematurely

Growing evidence suggests that precocious cell differentiation contributes to pathology in KS1. Previously, we found that both KMT2D-deficient neurons, another disease-relevant cellular model (Carosso et al., 2019), and chondrocytes display accelerated differentiation (Fahrner et al., 2019). For this study, we used our previously established ATDC5 chondrogenic cell lines (Fahrner et al., 2019), in which *Kmt2d* has been knocked out by CRISPR-Cas9 genome editing (*Kmt2d*^-/-^). Upon the addition of insulin on day 0 to induce their differentiation, ATDC5 cells undergo a chondrogenic differentiation program similar to that observed *in vivo* (Newton et al., 2012). Secretion of the glycosaminoglycan (GAG)-rich cartilage extracellular matrix (ECM) during the differentiation process was monitored by Alcian blue staining. Consistent with a precocious differentiation phenotype, *Kmt2d*^-/-^ chondrogenic cells secreted approximately three times more ECM than wild-type (*Kmt2d^+^*^/+^) cells already by day 7 of differentiation (**Figure 1A and 1B**). To gain a deeper understanding of the molecular mechanisms that trigger the premature cell differentiation in the absence of KMT2D, we performed single-cell RNA sequencing (sc-RNA-Seq) at four different days of differentiation (days 0, 4, 7, and 14). Visualization of the obtained sc-RNA-Seq data from the four timepoints using Uniform Manifold Approximation Projection (UMAP) revealed distinct clustering of the combined *Kmt2d*^+/+^ and *Kmt2d*^-/-^ cell population. A total of ten different cell subpopulations/clusters could be identified with clear differences in the number of cells within each cluster between the two genotypes. Induction of the differentiation process led to a shift on the UMAP from left to right and specifically an increase in cluster 1 (**Figure 1**). Closer inspection of the gene expression profiles of all clusters revealed that cells within cluster 1 likely represent pre-hypertrophic chondrocytes (**Supplementary Figure 1**) given their high expression of *Pth1r* (Chu et al., 2023), a parathyroid hormone receptor which is mainly expressed in the pre-hypertrophic zone of the growth plate (**Supplementary Figure 2A**). Notably, an increase in this cluster was more evident at earlier timepoints in *Kmt2d*^-/-^ cells than in wild-type cells (**Supplementary Figure 2B**), further supporting the notion that KMT2D deficiency drives accelerated differentiation and an early accumulation of pre-hypertrophic and hypertrophic cells.

**Figure 1:**
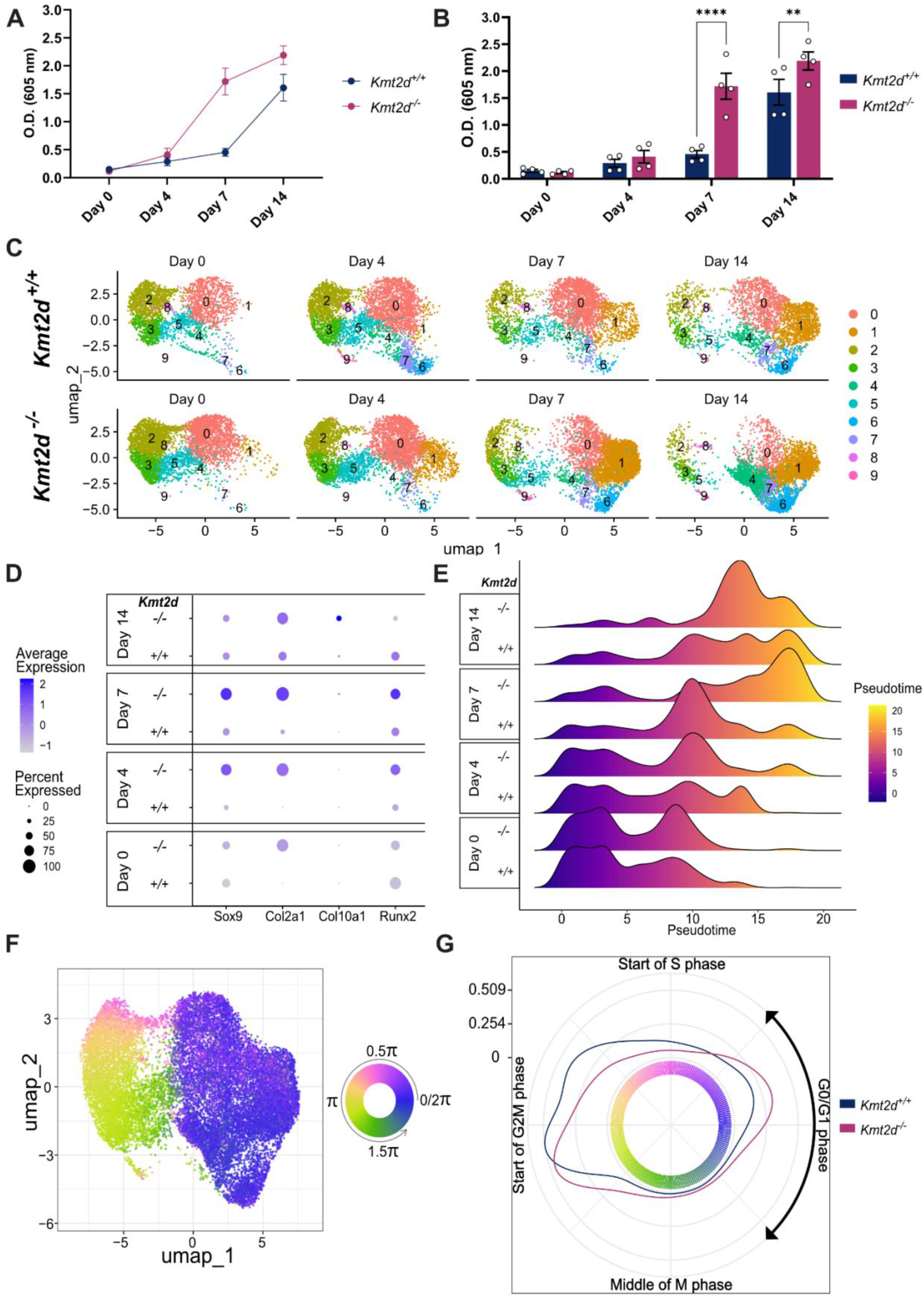
Precocious differentiation and cell cycle exit occurs early in KMT2D-deficient chondrocytes. A) Alcian blue staining and B) quantification of the cartilage extracellular matrix (ECM) secreted by *Kmt2d*^+/+^ and *Kmt2d*^-/-^ ATDC5 chondrogenic cells during the differentiation process (n=4). C) UMAP includes merging of both genotypes at multiple timepoints followed by harmony correction and cluster analysis. D) Chondrocyte marker gene expression throughout differentiation. The color scale represents expression levels and the dot size, the percentage of cells expressing the respective marker. E) Trajectory analysis with slingshot visualized as a density plot for cell count per pseudotime. F) UMAP projection of *Kmt2d^+/+^* cells colored by cell cycle state where 0.5π indicates the start of the S phase, π the start of the G2M phase, 1.5π the mid-M phase, and 1.75-0.25π the G1/G0 phase. G) Circular density plot of cell cycle state for day 0, estimated with the von Mises distribution and color-coded by phenotype. Data shown as mean ± SEM and significance calculated with two-way ANOVA. *P ≤ 0.05, **P ≤ 0.01.

Prior analyses of chondrocyte marker expression by quantitative real-time PCR (RT-qPCR) showed that an undifferentiated KMT2D-deficient ATDC5 cell population exhibited increased *Sox9* and *Col2a1* expression levels compared to wild-type cells (Fahrner et al., 2019). To determine whether this increase was due to a higher percentage of differentiated cells within the cell population or an overall increase in gene expression intensity, we examined the sc-RNA-Seq data for the master regulator of chondrocyte differentiation, *Sox9*. While the percentage of cells expressing *Sox9* remained similar between the two genotypes, we observed an increase in the average expression at day 0 of differentiation. However, during the differentiation process, both the percentage of cells expressing *Sox9,* and the *Sox9* expression level increased in *Kmt2d^-/-^* cells compared to *Kmt2d^+/+^* cells **(Figure 1D)**. For *Col2a1*, we observed increases in both the percentage of cells expressing these markers and their expression intensity from the beginning and throughout differentiation **(Figure 1D)**. On day 7 of differentiation, *Kmt2d^-/-^* cells began to express increasing levels of the hypertrophic marker C*ol10a1*, while the osteoblast marker *Runx2* could already be detected as early as four days after the induction of differentiation **(Figure 1D)**. To further explore the differentiation trajectory, we performed slingshot analysis on the sc-RNA-Seq data, followed by pseudotime-based trajectory analysis. Pseudotime was assigned to each cell, with higher values indicating greater differentiation **(Figure 1E)**. Consistent with earlier secretion of ECM **(Figure 1A and 1B)**, we detected an increased number of cells with a higher pseudotime within the *Kmt2d^-/-^* cell population **(Figure 1E)**, indicating an early tendency towards differentiation, which was evident according to Kolmogorov–Smirnov test from day 0 (P value = 8.216e-15) onwards.

Since the proliferation of chondrocytes in the growth plate determines longitudinal bone growth and the deposited cartilage extracellular matrix provides a scaffold for future bone formation (Stegen et al., 2019), early loss of their proliferative ability would inevitably impair successful bone elongation. Previously, we found that iPSC-derived neural stem/progenitor cells (NSPCs) from KS1 individuals exhibit a cell cycle defect that affects their proliferation (Carosso et al., 2019). To investigate whether *Kmt2d^-/-^* chondrocytes display similar changes in the cell cycle and differences in proliferative ability compared to *Kmt2d^+/+^* cells, we used the Tricycle package to predict the cell cycle stage of each individual cell. This analysis revealed a distinct separation between cycling (pink, yellow or green) and non-cycling cells (blue) on the UMAP **(Figure 1F)**. Interestingly, when quantifying cells in each phase, *Kmt2d^-/-^* cells exhibited a higher number of undifferentiated cells in the resting/non-proliferative G1/G0 phase compared to *Kmt2d^+/+^* cells **(Figure 1G)**. To experimentally validate the lower proliferation rate of *Kmt2d^-/-^* cells, we additionally performed staining of the undifferentiated *Kmt2d^+/+^ and Kmt2d^-/-^* cell populations with the dilutable marker Carboxyfluorescein succinimidyl ester (CFSE) and found that *Kmt2d^-/-^* cells divided less frequently than their wild-type counterparts **(Supplementary Figure 2C)**. These findings suggest that the earlier transition of KMT2D-deficient chondrocytes to a hypertrophic state is accompanied by a loss of proliferative ability – a process that is reminiscent of the differentiation of columnar chondrocytes into postmitotic hypertrophic chondrocytes during endochondral bone formation *in vivo* (Blumer, 2021).

### Dysregulated hypoxic response in KMT2D-deficient chondrocytes does not drive their premature hypertrophy

Next, we aimed to identify the biological pathways, the dysregulation of which might have caused the early entry of KMT2D-deficient chondrogenic cells into the chondrocyte differentiation program. For this purpose, we re-analyzed our previously published bulk RNA sequencing dataset from the same cell lines (Fahrner et al., 2019), identified the significantly up–, and downregulated genes **(Supplementary Table 1),** and performed gene set enrichment analysis (GSEA). Unexpectedly, the top enriched upregulated gene sets included “hypoxia” and “glycolysis” **(Figure 2A),** both of which are thought to play a role in the maintenance of the chondrocyte proliferative state and minimize chondrocyte hypertrophy *in vivo* (Hollander et al., 2022; Schipani et al., 2001). Hypoxic conditions in the proliferative zone of the epiphyseal growth plate necessitate the activation of the hypoxia response to allow the metabolic adaptation to a low oxygen tension and thus cellular survival (Chatzi et al., 2015; Schipani et al., 2001). The transition of proliferating chondrocytes to the hypertrophic state, however, requires a progressive downregulation of the hypoxic response and a bioenergetic reprogramming from glycolysis to oxidative phosphorylation to meet the increasing energy demands during maturation and prevent energy distress (Stegen et al., 2019). The observed upregulation of genes related to hypoxia and glycolysis in undifferentiated KMT2D-deficient cells **(Figure 2A)** made us thus wonder whether the dysregulation of normal chondrocyte maturation in the growth plate might be due to an impaired cellular response to the increasing oxygen level as they progress from the highly hypoxic proliferative to the less hypoxic hypertrophic zone (Chatzi et al., 2015; Schipani et al., 2001). To test this hypothesis, we first exposed the *Kmt2d^+/+^* and *Kmt2d^-/-^* chondrogenic cell populations to a supraphysiological oxygen level, i.e., 20% (hereafter referred to as “normoxia”), and monitored the expression of *Hif1a*, the master regulator of the hypoxic response, during the differentiation process. Interestingly, even under these non-hypoxic conditions, *Hif1a* expression levels remained elevated in *Kmt2d^-/-^* cells - not only before differentiation but also throughout the process **(Figure 2B)**. Consistent with anaerobic glycolysis being the primary energy-generating pathway regulated by HIF-1α, *Kmt2d^-/-^* cells exhibited significantly increased lactate secretion compared to *Kmt2d^+/+^* cells - 1.3-fold at day 4 and even 3-fold at day 7 of differentiation **(Figure 2C)**. The observed upregulation of HIF-1α signaling in *Kmt2d^-/-^* cells was rather unexpected, given that an active hypoxia response is associated with the suppression of chondrocyte hypertrophy (Hollander et al., 2022; Schipani et al., 2001). To directly assess whether increased HIF-1α signaling in KMT2D-deficient chondrogenic cells plays any role in their precocious differentiation, we knocked down *Hif1a* by RNA interference using two distinct siRNAs (siHif1a_1 and siHif1a_2) 12 hours before the induction of differentiation and monitored ECM secretion throughout the differentiation process. The *Hif1a* knockdown efficiency was evaluated by RT-qPCR at various days during differentiation. The average *Hif1a* transcript level in siHif1a_1 and siHif1a_2-treated *Kmt2d^-/-^* cells dropped to about 10% within the first 12 hours of siRNA treatment relative to those treated with non-targeting control siRNA, before it progressively increased again to approximately 50-60% over the 14-day differentiation period **(Supplementary Figure 3A)**. Quantification of ECM secretion during the differentiation process, however, did not reveal any significant differences between *Kmt2d^-/-^* chondrocytes treated with either *Hif1a-*targeting or non-targeting control siRNA **(Figure 2D)** despite their differences in *Hif1a* expression, strongly suggesting that the aberrant HIF-1α signaling in KMT2D-deficient chondrogenic cells is not causative for their precocious differentiation. To further confirm that activation of the hypoxia response *per se* does not increase the chondrocyte differentiation rate, we reduced the environmental oxygen level from 20% to 5% before we induced the differentiation of *Kmt2d^+/+^* cells. As expected, oxygen deprivation (5% O_2_) led to an increased anaerobic glycolysis rate, as evidenced by the elevated secretion of lactate **(Supplementary Figure 3B)**. Yet, we did not observe premature differentiation of the *Kmt2d^+/+^* chondrogenic cells in response to various levels of hypoxia **(Figure 2E and Supplementary Figure 3C)**, supporting our previous notion that the aberrant activation of the hypoxia response in KMT2D-deficient cells is likely not the responsible mechanism that underlies their precocious differentiation phenotype.

**Figure 2:**
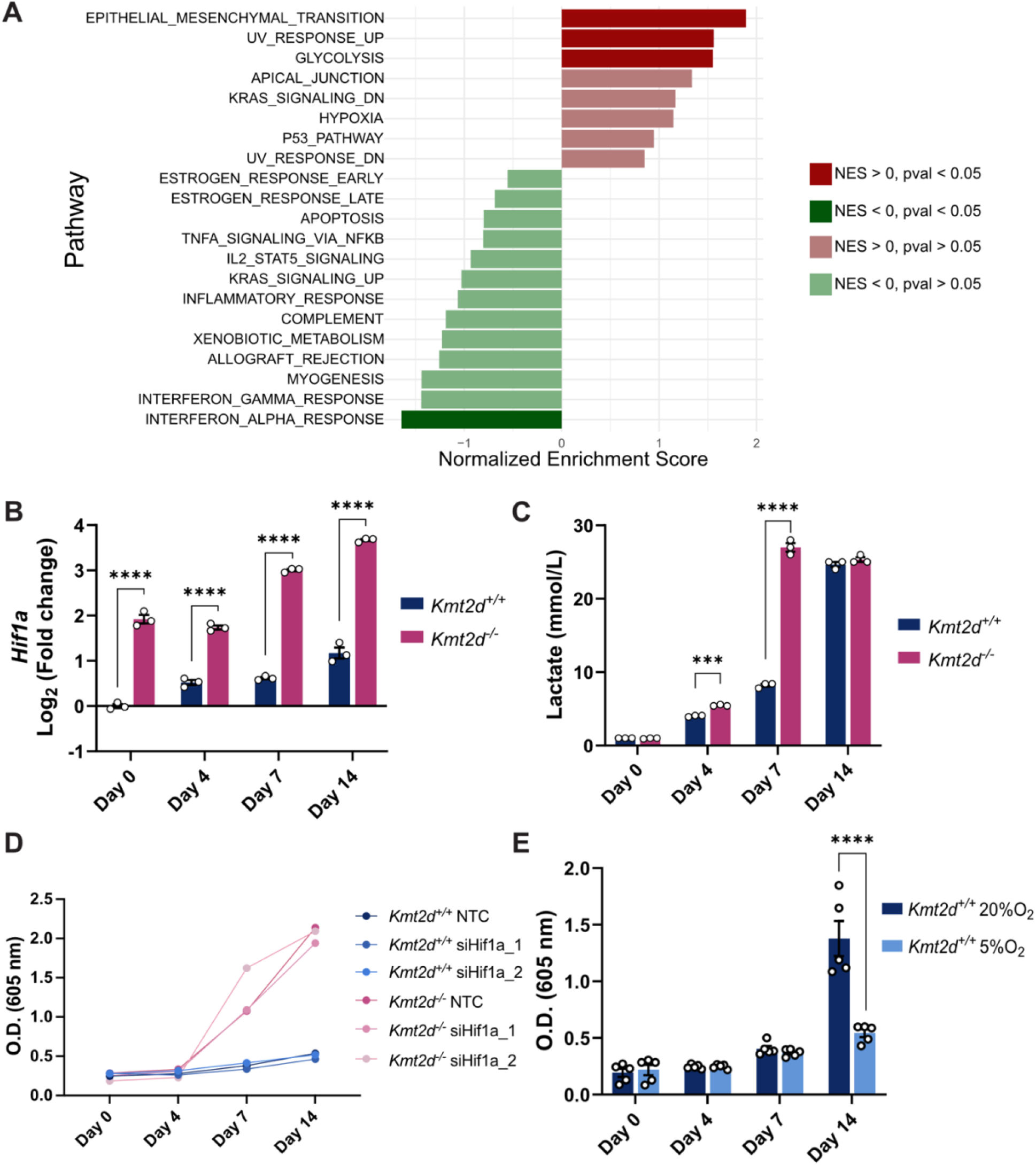
Hypoxia responses and glycolysis are enhanced in KMT2D-deficient chondrocytes, but are not the primary drivers of their precocious differentiation. A) Gene set enrichment analysis of differentially expressed genes in undifferentiated cells showing the top dysregulated pathways. Normalized Enrichment Score (NES) > 0 indicates upregulated, and NES < 0 downregulated pathways. B) *Hif1a* expression in *Kmt2d^+/+^* and *Kmt2d^-/-^* cells over the course of differentiation. C) Amperometry measurements for lactate in mmol/L from culture media at four timepoints during differentiation. D) Representative Alcian blue staining for siRNA-mediated knockdown of *Hif1a* in *Kmt2d^+/+^* and *Kmt2d^-/-^* cells. E) Quantification of Alcian blue staining for *Kmt2d^+/+^* cells differentiated in the presence of 20% and 5% environmental O_2_. Data shown as mean ± SEM. Statistical tests were performed with unpaired Student’s t-tests within individual timepoints (B, C) or two-way ANOVA (E). *P ≤ 0.05, **P ≤ 0.01, ***P ≤ 0.001, ****P ≤ 0.0001.

### KMT2D deficiency leads to increased mitochondrial ROS generation which drives early cellular senescence

Based on our finding that the aberrant upregulation of HIF-1α signaling and glycolysis under normoxia is not primarily responsible for the premature differentiation of KMT2D-deficient chondrocytes, we suspected that it may be a compensatory response to fulfill the increasing energy demand during chondrocyte maturation and prevent energy distress (Stegen et al., 2019). In fact, previous studies in fibroblasts from KS1 patients revealed deficiencies in the mitochondrial electron transport chain (ETC), leading to impaired mitochondrial respiration and glucose oxidation (Pacelli et al., 2020). The activation of the hypoxic response may thus help redirect cellular metabolism away from respiration and compensate for the loss of mitochondrial ATP production through anaerobic glycolysis.

In the normal epiphyseal growth plate, the progression from proliferating to pre-hypertrophic and hypertrophic chondrocytes is accompanied by an increase in oxidative phosphorylation (OXPHOS) and oxygen consumption rate (OCR). Given that undifferentiated *Kmt2d^-/-^* chondrogenic cells transcriptionally resembled an overall more differentiated state **(Figure 1C and 1E; Supplementary Figure 1)**, we expected the cells to show an increased OCR compared to *Kmt2d^+/+^* cells if their mitochondria were fully functional. To test whether *Kmt2d^-/-^* chondrogenic cells also exhibit a defect in mitochondrial respiration/OXPHOS, we measured the OCR under normoxic conditions using the Seahorse XF Cell Mito Stress Test. After measuring basal OCR at the beginning of the experiment, the subsequent addition of oligomycin, which inhibits ATP synthase (Complex V) in the ETC, is expected to decrease OCR, enabling calculation of mitochondrial ATP production. FCCP, an uncoupler of the ETC, is anticipated to maximize mitochondrial OCR, from which spare respiratory capacity can be determined. Finally, rotenone and antimycin A (R/A), inhibitors of complexes I and III of the ETC, respectively, completely block mitochondrial respiration, eliminating any OCR component derived from mitochondria. The remaining OCR after R/A treatment represents non-mitochondrial respiration. We measured the OCR in two independent biological replicates (Rep. 1, Rep 2) for both *Kmt2d^+/+^* and *Kmt2d*^-/-^ cells, with data from each replicate presented separately **(Rep. 1, Figure 3A and 3B; Rep. 2, Supplementary Figure 4A-B)**. Surprisingly, quantification of the OCR showed that the basal respiration (P value ≤ 0.0001 for Rep.1 and Rep. 2), maximal respiration after FCCP injection (P value ≤ 0.0001 for Rep.1 and Rep. 2) and the spare respiratory capacity (P value ≤ 0.0001 for Rep. 1, P value = 0.0032 for Rep. 2) were decreased in *Kmt2d^-/-^* cells compared to *Kmt2d^+/+^* cells **(Figure 3A and 3B, Supplementary Figure 4A-B)**. These findings indicate a reduced capacity for mitochondrial respiration in energy production, potentially explaining the need to activate the hypoxic response, especially as energy demands increase during chondrocyte maturation. A reduced respiration rate can arise either from intrinsic defects in the ETC or from an overall decrease in the number of functional mitochondria per cell. To distinguish between these possibilities, cells can be cultured in galactose-containing medium, which forces reliance on OXPHOS for ATP production independent of mitochondrial abundance. To assess whether *Kmt2d^-/-^*cells retain functional mitochondria, we replaced glucose-rich medium with medium containing 10 mM galactose as the sole carbon source for 1 h prior to repeating the Seahorse assay. Interestingly, while *Kmt2d^+/+^* cells increased their basal respiratory rate by approximately 13.5% **(Figure 3C and 3D, *top panel*)**, *Kmt2d^-/-^* cells failed to significantly upregulate their basal mitochondrial respiration despite the lack of glucose in the cell culture medium **(Figure 3C and 3D, *bottom panel*)**. These findings indicate that *Kmt2d^-/-^* cells are unable to efficiently utilize the mitochondrial ETC for energy production due to an underlying defect in the ETC. In our bulk RNA-seq dataset from undifferentiated chondrocytes, we identified 90 nuclear- and mitochondrial-encoded genes that give rise to ETC complex subunits. Analysis of their expression levels showed that the majority were not significantly altered in *Kmt2d*^-/-^ cells, with only about ∼27% (24/90 genes) displaying significant differential expression **(Supplementary Figure 5 and Supplementary Table 1)**. Importantly, while 75% (18/24) of differentially expressed ETC genes were downregulated, 25% (6/24) were upregulated, demonstrating a bidirectional change in expression rather than uniform downregulation. These genes were distributed across all ETC complexes and did not exhibit consistent directionality within individual complexes **(Supplementary Figure 5)**. Together, these results suggest that the observed ETC defect is unlikely to be explained solely by transcriptional changes caused by loss of KMT2D. Defects in the ETC often lead to increased reactive oxygen species (ROS) production (Pacelli et al., 2020). While hypertrophic growth plate chondrocytes naturally exhibit higher ROS levels due to their proximity to oxygen-enriched blood vessels and utilization of OXPHOS (Hollander et al., 2022), pathophysiological ROS levels can prematurely inhibit chondrocyte proliferation and stimulate chondrocyte hypertrophy (Morita et al., 2007). To test whether the observed mitochondrial dysfunction in *Kmt2d^-/-^* chondrogenic cells increases the generation of superoxide (•O_2_^-^), the first ROS produced by mitochondrial ETC, we stained the undifferentiated *Kmt2d^-/-^* and control *Kmt2d^+/+^* cells with the fluorescent superoxide-specific probe MitoSox (Kauffman et al., 2016) and analyzed their fluorescence intensity by flow cytometry **(Figure 3E)**. Indeed, we detected an approx. 1.5-fold increased superoxide production by *Kmt2d^-/-^* cells compared to their wild-type counterparts. Given that superoxide is rapidly converted to hydrogen peroxide (H_2_O_2_), one of the most prevalent ROS in the cell (Ulfig & Jakob, 2024), we next assessed the intracellular H_2_O_2_ level using 2’-7’-dichlorodihydrofluorescein diacetate (DCFDA), a fluorogenic probe that is oxidized mainly by H_2_O_2_, leading to green fluorescence (Eruslanov & Kusmartsev, 2010). Surprisingly, DCFDA staining revealed a 56% decrease in H_2_O_2_ accumulation in undifferentiated *Kmt2d^-/-^* vs. *Kmt2d^+/+^* cells **(Figure 3F)**. In an attempt to resolve this apparent paradox, we examined the expression level of the *Nfe2l2* gene encoding NRF2, the master regulator of the cellular antioxidant response, which is activated by the disruption of mitochondrial redox homeostasis and induces the expression of ROS-neutralizing antioxidant enzymes (Cvetko et al., 2021). In fact, we found increased expression of *Nfe2l2* in *Kmt2d^-/-^* cells in our sc-RNA-Seq dataset both at day 0 and day 4 of differentiation **(Supplementary Figure 4C),** and in the re-analyzed, previously published bulk-RNA-Seq dataset from undifferentiated cells (Fahrner et al., 2019) **(Supplementary Table 1;** Log2FoldChange=0.54, padj=5e-3). More importantly, *Kmt2d^-/-^* cells show increased expression of the NAD(P)H quinone oxidoreductase 1 (*Nqo1*) **(Supplementary Table 1;** Log2FoldChange=1.2, padj=0.039), a key gene activated by NRF2 in response to oxidative stress, confirming a higher NRF2 activity in KMT2D-deficient cells that may be responsible for the overall decreased intracellular H_2_O_2_ levels despite elevated mitochondrial •O_2_^-^production. As mentioned earlier, maturing chondrocytes naturally increase their utilization of OXPHOS for energy generation and hence, ROS production (Hollander et al., 2022). We thus wondered whether the observed ETC dysfunction in *Kmt2d^-/-^* cells may exacerbate ROS production during this process, leading to oxidative distress – a harmful cellular state which rapidly promotes cellular senescence and cell death (Morita et al., 2007). Strikingly, measurement of H_2_O_2_ accumulation using DCFDA during the differentiation of *Kmt2d^-/-^* and *Kmt2d^+/+^* chondrocytes revealed approximately 9-fold higher levels in KMT2D-deficient cells at day 7 of differentiation **(Figure 3F)**. This drastic increase in cellular ROS correlated with increased ECM secretion **(Figure 1B)** and hypertrophic marker expression **(Figure 1D)**, supporting the direct link between ROS accumulation and the induction of chondrocyte hypertrophy (Morita et al., 2007).

**Figure 3:**
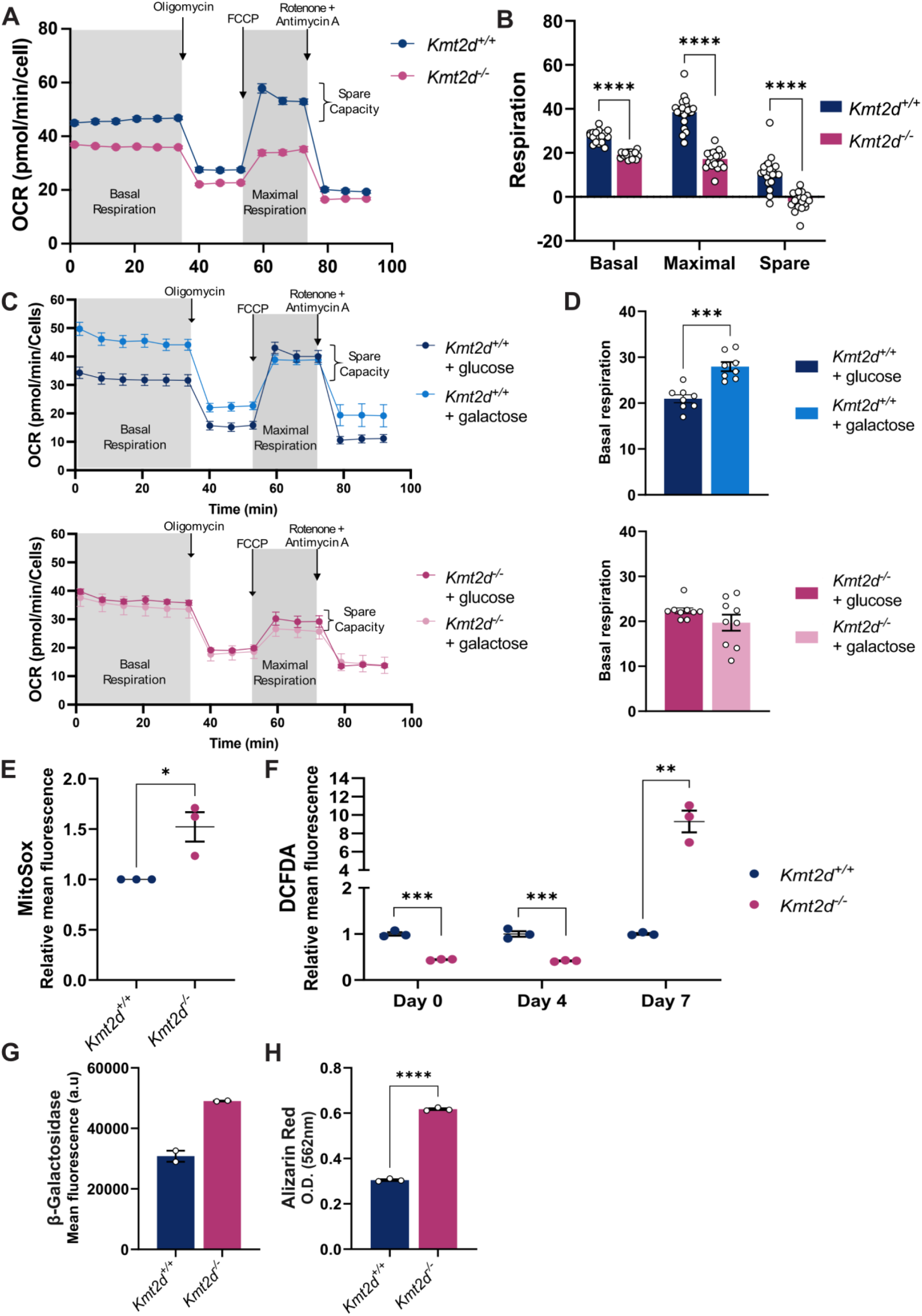
Impaired mitochondrial respiration causes increased ROS production and early cellular senescence. Oxygen consumption rate (OCR) of undifferentiated *Kmt2d^+/+^* and *Kmt2d^-/-^* cells measured with a Seahorse assay. A) Average respiration rate across all technical replicates over time. B) Mitochondria-specific basal and maximal respiration were calculated by subtracting the non-mitochondrial respiration from the total basal and maximal respiration, respectively. Spare respiration was calculated by subtracting the mitochondria-specific basal respiration from the mitochondria-specific maximal respiration. C) Average respiration rate over time and D) quantification of basal mitochondria-specific respiration upon incubation of *Kmt2d^+/+^* and *Kmt2d^-/-^* cells with glucose or galactose as carbon source for energy production. E) Mitochondrial ROS were quantified with the superoxide-specific probe MitoSox in undifferentiated cells. F) Intracellular H_2_O_2_ was detected with DCFDA on days 0, 4, and 7 of differentiation. G) Analysis of β-galactosidase activity in *Kmt2d^+/+^* and *Kmt2d^-/-^*cells at day 20 of differentiation using a fluorescence-based assay. H) Quantification of Alizarin red staining on day 30 of differentiation. Data shown as mean ± SEM. Statistical tests were performed with unpaired Student’s t-tests. *P ≤ 0.05, **P ≤ 0.01, ***P ≤ 0.001, ****P ≤ 0.0001.

Given that excessive ROS production is known to induce cellular senescence, a hallmark of terminally differentiated hypertrophic chondrocytes (Bolduc et al., 2019; Morita et al., 2007), we next asked whether the drastic increase in ROS production during the accelerated maturation of KMT2D-deficient chondrocytes may trigger early/premature senescence of hypertrophic cells. Interestingly, examination of the cluster distribution in the sc-RNA-Seq dataset from day 14 of differentiation revealed an expansion of particular subpopulations in *Kmt2d^-/-^* cells, such as cluster 4, which expresses DNA damage-inducible factors (*Ddit3, Gadd45a* and *Ppp1r15a*), and cluster 6, which expresses mitochondrial genes **(Supplementary Figure 1)**. These gene expression profiles suggest that these cell populations may represent pre-apoptotic and apoptotic cells, further supporting the idea of a premature transition of *Kmt2d^-/-^* cells toward senescence. To directly test this, we measured the activity of senescence-associated β-galactosidase (β-gal) in *Kmt2d^+/+^* and *Kmt2d^-/-^* cells at day 20 of differentiation **(Figure 3G)**. β-Gal activity was nearly twice as high in *Kmt2d^-/-^* chondrocytes as in the wild-type control, strongly suggesting that hypertrophic *Kmt2d^-/-^* cells indeed undergo early cell senescence. Senescence of hypertrophic chondrocytes is known to promote ECM calcification (Cho et al., 2024). To test whether the early senescence of hypertrophic *Kmt2d^-/-^* cells causes premature matrix calcification, we stained the cells with Alizarin Red at day 30 of differentiation, a chemical compound that specifically binds to calcium (Puchtler et al., 1969). In fact, we observed a 2-fold increase in the staining intensity of KMT2D-deficient cells compared to wild-type cells **(Figure 3H)**, pointing towards an overall dysregulated endochondral ossification process.

### Targeting increased ROS accumulation rescues accelerated differentiation of KMT2D-deficient chondrogenic cells

Our results strongly suggested that excessive accumulation of ROS in KS1 contributes to the premature transition of proliferating chondrocytes toward hypertrophic and senescent states. We therefore suspected that decreasing ROS abundance during the differentiation of KMT2D-deficient cells could restore a normal differentiation rate and prevent premature hypertrophy and cellular senescence. Of note, previous studies on the role of ROS in chondrocyte differentiation revealed that a moderate increase in endogenous ROS is required for normal chondrogenic differentiation, as antioxidant treatments that prevent ROS accumulation completely inhibit this process (Ki et al., 2010; Morita et al., 2007). These differentiation-associated ROS are thought to be primarily derived from the NADPH oxidases NOX1, NOX2, and NOX4, which generate superoxide (O₂•⁻) by transferring electrons from NADPH to molecular oxygen (Ki et al., 2010; Morita et al., 2007). Importantly, defects in the mitochondrial ETC would be expected to generate additional ROS on top of NOX-derived oxidants, potentially pushing total ROS levels beyond a critical threshold that shifts redox signaling from a pro-differentiation cue to a driver of premature hypertrophy and senescence (Ki et al., 2010; Morita et al., 2007). To test this hypothesis, we used the mitochondria-targeted antioxidant mitoquinone (MitoQ), which was found to efficiently alleviate oxidative stress caused by aberrant mitochondrial •O_2_^-^ production in various cell types (Mao et al., 2022; Murphy, 2001), and pre-treated both *Kmt2d^+/+^* and *Kmt2d^-/-^* cells with either 100 or 200 µM MitoQ for 24 hours before we induced their differentiation. Treatment with MitoQ reduced the accumulation of cellular H₂O₂ only partially, likely because it selectively targets the mitochondria-derived fraction of total ROS rather than NOX-derived oxidants **(Figure 4A)**. Notably, despite this limited reduction in overall H₂O₂ levels, MitoQ treatment decreased the excessive ECM secretion by *Kmt2d^-/-^* cells at day 7 of differentiation in a dose-dependent manner **(Figure 4B)**, indicating that mitochondrial ROS contribute to the aberrant differentiation phenotype. In fact, *Kmt2d^-/-^* cells treated with the higher MitoQ dose, i.e., 200 µM, differentiated even at a similar rate as untreated *Kmt2d^+/+^* cells, as judged by the amount of deposited ECM by day 14 of differentiation **(Figure 4B)**, visualizing the direct correlation between the endogenous mitochondrial ROS level and rate of differentiation. While no significant changes were observed in early-chondrogenic marker gene expression, *Sox9* or *Col2a1* **(Supplementary Figure 4D-E)**, expression of the hypertrophic marker *Col10a1* was decreased in Kmt2d^-/-^ cells, consistent with slowed-down ECM deposition upon their treatment with MitoQ, pointing towards unaltered propensity to initiate differentiation but delayed hypertrophy later in the process **(Figure 4C).** Finally, treatment with MitoQ fully rescued the early cellular senescence of Kmt2d^-/-^ chondrocytes **(Figure 4D)**. Interestingly, the highest dose of MitoQ increased senescence in *Kmt2d^+/+^* cells, as indicated by elevated β-galactosidase staining **(Figure 4D)**, and completely blocked differentiation **(Figure 4B)**. This suggests that in cells with an intact ETC and no excess mitochondrial ROS, high-dose MitoQ may be detrimental, potentially by over-scavenging ROS and inducing reductive stress rather than restoring redox balance. An alternative strategy to alleviate oxidative stress is to lower environmental oxygen levels, which restricts oxygen availability within mitochondria and consequently reduces mitochondrial superoxide generation. Intriguingly, while the exposure of *Kmt2d^+/+^* cells to lower oxygen levels (5% vs. 20% O₂) had little effect on intracellular ROS at day 7 of differentiation, *Kmt2d^-/-^* cells not only showed a marked reduction in ROS accumulation **(Figure 4E)** but also slowed their differentiation rate **(Figure 4F)**, supporting the hypothesis that the total intracellular ROS level, which is dependent on oxygen availability, dictates the pace of the chondrogenic differentiation/maturation process.

**Figure 4:**
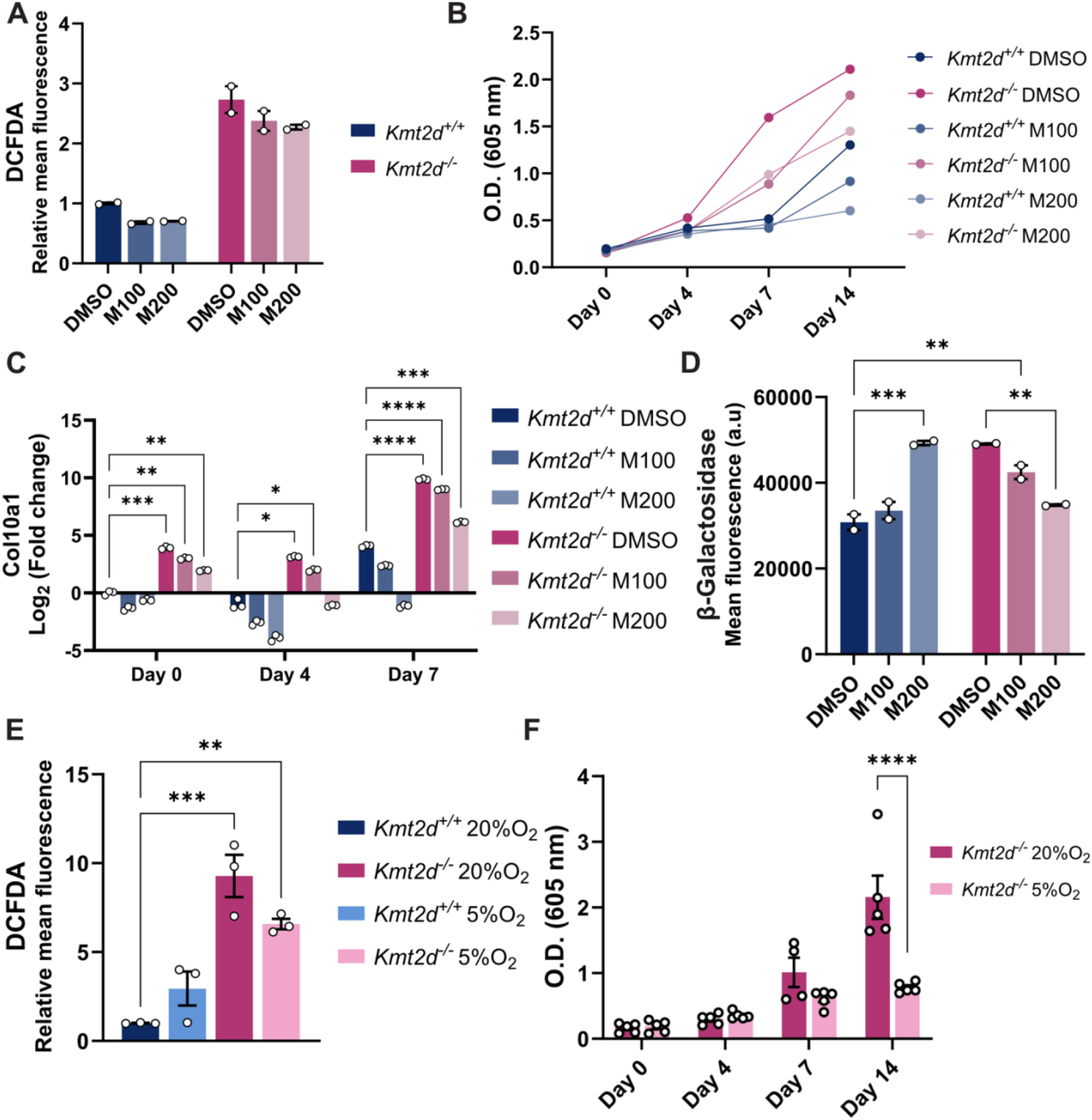
Targeting ROS and oxygen accumulation rescues precocious differentiation of KMT2D-deficient chondrogenic cells. A) Relative mean DCFDA fluorescence of *Kmt2d^+/+^* and *Kmt2d^-/-^* cells treated with different concentrations of MitoQ (M100: 100 µM; M200: 200 µM) or DMSO as control at day 7 of differentiation (n=2). B) Representative Alcian blue staining quantification of the deposited ECM. C) *Col10a1* expression of *Kmt2d^+/+^* and *Kmt2d^-/-^* cells with or without MitoQ treatment during differentiation. D) β-Galactosidase mean fluorescence intensity as the read-out for cellular senescence at day 20 of differentiation in the presence of the indicated concentrations of MitoQ or DMSO. E) Relative DCFDA fluorescence mean at day 7 of differentiation in the presence of either 20% or 5% environmental O_2_. F) Alcian blue staining for *Kmt2d^-/-^* cells differentiated in the presence of 20% and 5% environmental O_2_. Data shown as mean ± SEM. Statistical tests were performed with two-way ANOVA or mixed-effects ANOVA. *P ≤ 0.05, **P ≤ 0.01, ***P ≤ 0.001, ****P ≤ 0.0001.

### Minor increases in oxygen levels drive hypertrophy in the fetal growth plate

Our *in vitro* findings prompted us to ask whether altered oxygen availability and redox imbalance similarly affect chondrocyte maturation *in vivo*. To test this, we utilized our previously characterized mouse model for KS1 (*Kmt2d^+/βGeo^*), which displays a skeletal growth deficiency phenotype similar to that seen in individuals with KS1 (Fahrner et al., 2019). By late embryogenesis around E17.5, the growth plate of long bones has entered advanced stages of endochondral ossification. Proliferating chondrocytes have exited the cell cycle, begun terminal differentiation, and express markers of hypertrophy such as COLX, which are essential for longitudinal bone growth. Increased ROS levels have been observed specifically in pre-hypertrophic and hypertrophic chondrocytes in E17.5 embryos (Morita et al., 2007). We thus examined the femur growth plates from E17.5 *Kmt2d^+/+^* and *Kmt2d^+/βGeo^* embryos to assess whether pathological changes in growth plate organization resemble the oxidative stress–driven differentiation defects observed in the *in vitro* chondrocyte culture **(Figure 5)**. Given that an oxygen gradient exists across the growth plate (i.e., from the hypoxic proliferative zone to the oxygen-rich hypertrophic zone) (Schipani et al., 2001), we wondered whether increased oxygen exposure at the transition between the hypoxic proliferative zone and the more oxygen-rich hypertrophic region correlates with premature chondrocyte hypertrophy in *Kmt2d^+/βGeo^* embryos. In *Kmt2d^+/+^* embryos, staining with the hypoxia-labeling agent EF5 clearly demarcated the hypoxic core of the growth plate **(Figure 5A, turquoise line)**, while the COLX-positive hypertrophic zone **(Figure 5A, yellow line)** was largely devoid of a EF5 signal, consistent with its localization to a more oxygen-rich region adjacent to the blood vessels. Notably, analysis of distal femur growth plates revealed no significant differences in overall growth plate length **(Figure 5B)** or in the relative sizes of the hypoxic **(Figure 5C)** and hypertrophic zones **(Figure 5D)** between *Kmt2d^+/+^* and *Kmt2d^+/βGeo^* embryos. Compared to wild-type, *Kmt2d^+/βGeo^* embryos exhibited a significant increase in the overlap between EF5-positive hypoxic cells and the COLX-positive hypertrophic region **(Figure 5E)**. This overlap was most pronounced at the transition zone between the hypoxic core and the EF5-negative **(Figure 5A, orange line)**, oxygen-rich hypertrophic region, indicating that in *Kmt2d^+/βGeo^* growth plates, chondrocytes initiate hypertrophic differentiation already at lower oxygen levels than normal. These findings support the notion that even subtle increases in oxygen availability along the physiological gradient are sufficient to trigger premature hypertrophy. Collectively, our findings provide evidence that oxidative stress-induced early chondrocyte hypertrophy and senescence contribute to the disrupted endochondral ossification in KS1. These novel insights not only enhance our understanding of the mechanisms underlying growth deficiency in KS1 but also highlight the potential of therapeutic strategies specifically targeting ROS regulation.

**Figure 5:**
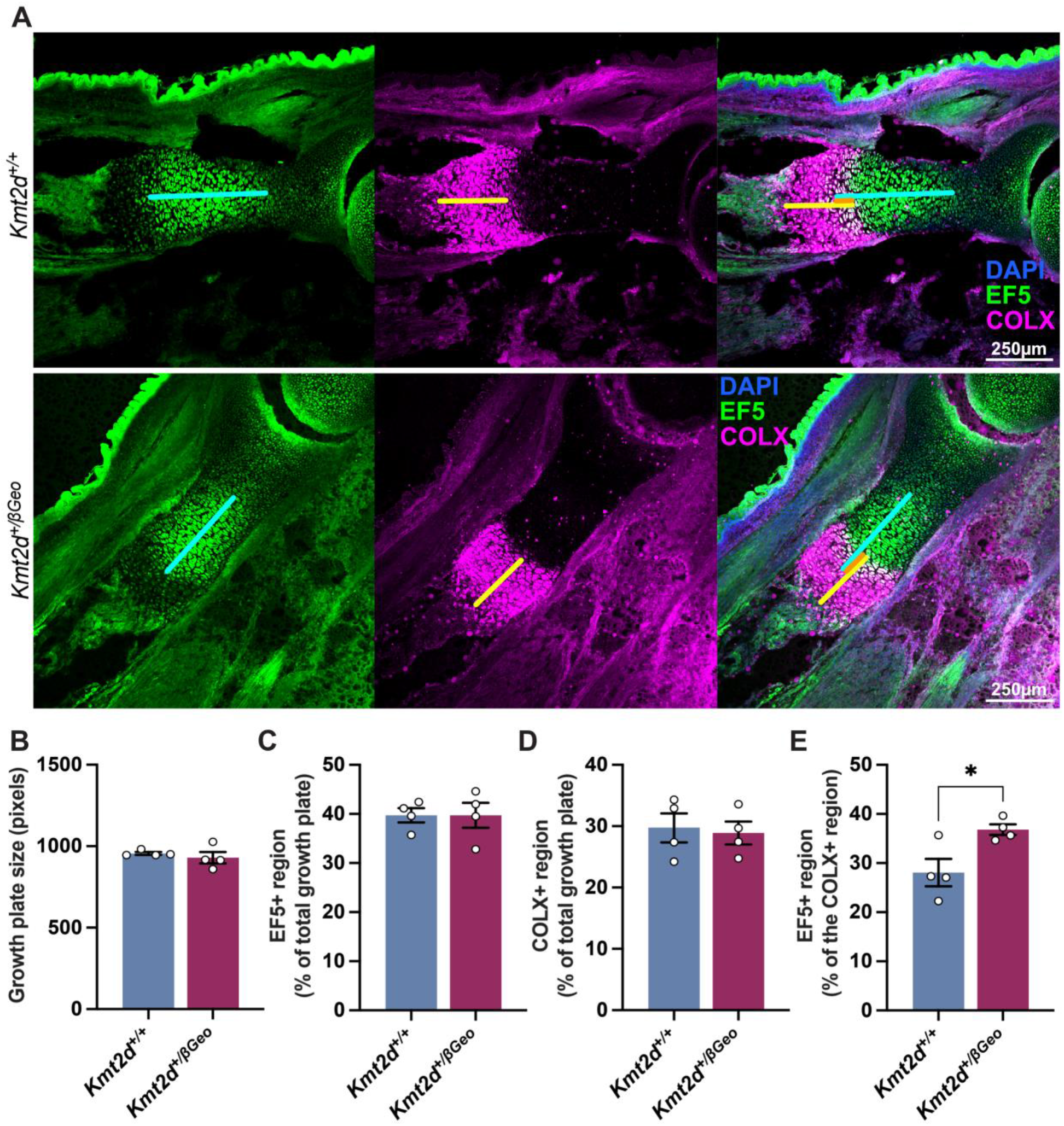
Minor changes in oxygen trigger hypertrophy in the E17.5 *Kmt2d^+/βGeo^* femur growth plate. A) Immunofluorescence staining of the distal femur growth plate of E17.5 *Kmt2d^+/+^* (top panel) and *Kmt2d^+/^*^β^*^Geo^* (bottom panel) embryos for hypoxia marker EF5 (green) and the hypertrophic marker collagen 10 (COLX; magenta). Nuclei were stained with DAPI (blue). Turquoise line indicates the length of the hypoxic region, yellow line the length of the hypertrophic region, and orange line the length of the overlap of the hypoxic and hypertrophic regions. Size of B) total growth plate shown in pixels, C) hypoxic (EF5+) region shown as the percentage of total growth plate, D) hypertrophic (COLX+) region shown as the percentage of total growth plate, and E) the overlap the hypoxic and hypertrophic region in the distal femur growth plate of E17.5 *Kmt2d^+/+^* and *Kmt2d^+/^*^β^*^Geo^* embryos defined as the proportion (in %) of the COLX+ region that is EF5+. Scale bar = 250 µm.

## Discussion

### Mitochondrial ROS generation during chondrocyte maturation – A delicate balance between the induction of normal hypertrophy and premature senescence

The process of endochondral ossification in the growth plate is highly sensitive to the timing of chondrocyte proliferation, hypertrophy, and senescence. We previously demonstrated that skeletal abnormalities and postnatal growth deficiency in KS1 patients likely result from accelerated chondrocyte differentiation, which disrupts the tightly coordinated sequence of endochondral ossification and ultimately leads to shortened long bones and craniofacial abnormalities (Fahrner et al., 2019). However, the mechanisms by which reduced KMT2D levels translate into impaired skeletal growth have remained unclear. Here, we show that KMT2D-deficient chondrocytes exhibit impaired mitochondrial OXPHOS, resulting in increased mitochondrial ROS production in an oxygen-dependent manner.

Although chondrocytes require a certain level of ROS for normal differentiation, these physiological ROS appear to be predominantly generated by NOX enzymes. Indeed, depletion of either NOX2 or NOX4 blocks both ROS production and chondrocyte differentiation *in vitro* (Ki et al., 2010). NOX-derived ROS seem particularly important during the early stages of differentiation, where they may initiate maturation and promote the stepwise transition from proliferating, primarily glycolytic chondrocytes to hypertrophic chondrocytes that increasingly rely on mitochondrial OXPHOS in addition to glycolysis to meet rising energy demands (Hollander et al., 2022). This suggests that mitochondria-derived ROS are not essential for initiating chondrocyte maturation but rather emerge as a byproduct of enhanced mitochondrial metabolism at later stages. Consistent with this notion, neutralization of mitochondrial ROS with the mitochondria-targeted antioxidant MitoQ prior to differentiation did not prevent the premature expression of maturation markers in KMT2D-deficient chondrocytes, nor did it reduce their propensity to differentiate prematurely.

Nevertheless, as mitochondrial metabolism increases during hypertrophic maturation (Hollander et al., 2022; Stegen et al., 2019), mitochondrial ROS inevitably accumulate and must be tightly controlled, as excessive ROS can promote premature senescence and cell death. This becomes particularly relevant *in vivo*, where the growth plate is organized along an oxygen gradient from the hypoxic proliferative zone to the more oxygen-rich hypertrophic region. This gradient influences mitochondrial ETC activity and, consequently, ROS generation. In line with this, Morita et al. showed that in the murine E17.5 growth plate, ROS levels are naturally elevated in pre-hypertrophic and hypertrophic chondrocytes exposed to higher oxygen levels compared to proliferating chondrocytes residing in the hypoxic zone. It is therefore plausible that disturbances in mitochondrial function or antioxidant defenses—which further increase ROS accumulation—would disproportionately affect cells in the oxygen-rich hypertrophic zone, where mitochondrial ROS production is already elevated. Supporting this concept, mice lacking the oxidative stress-responsive kinase ATM exhibit increased ROS levels in the growth plate and enhanced chondrocyte hypertrophy – a phenotype that can be rescued by antioxidant treatment with N-acetyl cysteine (Morita et al., 2007).

Similarly, in *Kmt2d^+/βGeo^* embryos, even modest increases in oxygen exposure at the transition between the hypoxic proliferative zone and the hypertrophic region were sufficient to trigger premature hypertrophy. The increased overlap between EF5-positive hypoxic areas and COLX-positive hypertrophic zones indicates that the oxygen/redox threshold required to initiate hypertrophy is lowered in the absence of KMT2D, likely due to elevated basal ROS levels in these cells. Comparable principles have been described in other tissues, where progenitor cell fate decisions are highly sensitive to subtle changes in ROS or oxygen availability (Chandel, 2014; Hamanaka & Chandel, 2010). Notably, KS1 patient-derived fibroblasts and neurons also display ETC defects and increased ROS, suggesting that redox-sensitive differentiation defects may represent a broader, unifying mechanism underlying multiple manifestations of the syndrome (Carosso et al., 2019; Pacelli et al., 2020). Taken together, these observations support the idea that mitochondrial OXPHOS dysfunction and untimely increases in ROS act as key drivers of early hypertrophy and premature senescence in KMT2D-deficient chondrocytes.

This model is biologically plausible, as other mitochondrial respiratory chain disorders are likewise associated with growth restriction and short stature (Holzer et al., 2019). Consistently, mice with impaired respiratory chain function in cartilage develop postnatal growth deficiency and premature growth plate closure (Holzer et al., 2019). We therefore propose that excess mitochondrial ROS production beyond physiological levels, caused by ETC dysfunction and exacerbated by increased oxygen availability near the primary spongiosa, accelerates chondrocyte differentiation along the oxygen gradient and induces early cellular senescence in KS1 and potentially in related neurocristopathies.

Interestingly, despite the accelerated differentiation observed *in vitro*, we did not detect significant differences in the relative sizes of distinct growth plate zones between E17.5 *Kmt2d^+/+^* and *Kmt2d^+/βGeo^* embryos. This initially appeared counterintuitive. However, recent evidence suggests that the metabolic shift from glycolysis to mitochondrial OXPHOS in hypertrophic chondrocytes occurs predominantly after birth (Holzer et al., 2019), implying that mitochondrial respiration—and thus mitochondria-derived ROS—may not yet be fully engaged at E17.5. In addition, growth deficiency in KS1 patients becomes evident primarily postnatally (Schott et al., 2016), with intrauterine growth largely preserved until late gestation. Together, these observations suggest that premature senescence or cell loss of hypertrophic chondrocytes *in vivo* may manifest predominantly after the postnatal glycolysis-to-OXPHOS transition, potentially triggered by increased oxygen supply to the growth plate after birth (Holzer et al., 2019). In line with this interpretation, we previously detected differences in growth plate architecture in 6-week- and 4.5-month-old *Kmt2d^+/βGeo^* mice (Fahrner et al., 2019).

However, since oxygen levels within distinct growth plate zones have not yet been directly quantified across developmental and postnatal stages, the precise relationship between oxygen availability, mitochondrial ROS, hypertrophy, and senescence remains to be defined. Future studies should therefore address these parameters over time to understand how mitochondrial dysfunction influences growth plate structure and long bone elongation in KS1.

### Mitochondrial dysfunction in KMT2D-deficient chondrocytes: Cause or consequence of oxidative stress?

One central question that remains is how KMT2D deficiency mechanistically leads to mitochondrial dysfunction and increased ROS production in chondrocytes. Given that KMT2D is a histone methyltransferase that deposits the transcription-activating marks H3K4me1 at enhancers and H3K4me3 at promoters across a broad range of genes, one plausible explanation is that its loss alters the methylation landscape of genes involved in mitochondrial processes, such as mitochondrial biogenesis (e.g., PGC-1α) or mitophagy. Disruption of these pathways could reduce the number of functional mitochondria and promote the accumulation of damaged mitochondria, with the latter contributing to increased ROS generation. However, when we forced *Kmt2d^+/+^* and *Kmt2d^-/-^*chondrocytes to rely on mitochondrial respiration by culturing them in galactose-containing medium, *Kmt2d^-/-^* cells failed to increase their basal respiration rate. Under these conditions, cells must depend on OXPHOS for ATP production, and healthy mitochondria normally respond with enhanced respiratory activity. The inability of *Kmt2d^-/-^*chondrocytes to do so indicates a functional defect in their respiratory machinery, rather than merely a reduced mitochondrial number, suggesting that the mitochondria present are intrinsically compromised in their capacity to generate energy efficiently. Along these lines, Pacelli et al. reported that KMT2D-deficient mouse embryonic fibroblasts (MEFs) display reduced protein levels of ETC complexes I and IV compared to wild-type cells, accompanied by decreased OCR, lower mitochondrial membrane potential, and increased ROS production (Pacelli et al., 2020). At first glance, the reduced abundance of ETC complexes could suggest that loss of KMT2D directly alters the transcription of ETC components. However, we did not detect significant differences in the mRNA expression of most nuclear- or mitochondrial-encoded ETC subunits in KMT2D-deficient compared to wild-type chondrocytes. Although a few genes were differentially expressed, their functional impact on complex assembly remains to be determined. Overall, this implies that the decreased protein levels of ETC complexes are unlikely to be primarily driven by transcriptional changes.

Intriguingly, mitochondrial DNA resides in close proximity to the ETC and is therefore particularly vulnerable to oxidative damage. Excess ROS can induce mutations or lesions in mitochondrial DNA, and thus in mitochondria-encoded ETC subunits (Cline, 2012). Even subtle defects in these core subunits can disrupt proper complex assembly, leading to instability and subsequent degradation of the entire respiratory complex. In this scenario, elevated ROS would precede and drive ETC dysfunction, with reduced complex abundance representing a downstream consequence of oxidative mitochondrial DNA damage rather than a direct effect of altered chromatin regulation.

This, however, raises the question of what drives the initial increase in ROS. One possibility is a reduced antioxidant capacity in KMT2D-deficient cells. Indeed, in prostate cancer, loss of KMT2D was shown to decrease enhancer-associated H3K4me1 and H3K27ac levels, impair FOXO3 binding, suppress transcription of antioxidant genes, and thereby promote ROS accumulation and DNA damage (Lv et al., 2019). In this scenario, defective antioxidant gene expression could act as the primary trigger for ROS elevation, which in turn compromises mitochondrial function.

However, our data argue against a complete breakdown of antioxidant defenses in KMT2D-deficient chondrocytes. We observed upregulation of NRF2, the master regulator of the cellular antioxidant response, and many NRF2 target genes overlap functionally with FOXO3-dependent antioxidant pathways. This suggests that antioxidant capacity is not entirely lost, although it may be insufficient to counterbalance increased ROS production. Thus, it remains to be determined whether elevated ROS levels are the primary cause of ETC dysfunction or whether subtle intrinsic defects in the respiratory chain initiate ROS accumulation, establishing a feed-forward cycle.

Although the precise sequence of events linking KMT2D deficiency to mitochondrial dysfunction and ROS accumulation remains unresolved, our data demonstrate that limiting excess ROS restores a more physiological chondrogenic differentiation rate and prevents premature senescence. This identifies mitochondrial ROS as a potential therapeutic target, particularly in the postnatal period when the switch toward increased mitochondrial respiration renders hypertrophic chondrocytes more susceptible to oxidative stress. Notably, antioxidant treatment has shown beneficial effects in mouse models of other growth and mitochondrial disorders, including pseudoachondroplasia (PSACH), diastrophic dysplasia, and Leigh syndrome (Bakare et al., 2021; Monti et al., 2015; Posey et al., 2015), where reducing oxidative stress alleviated pathological features to varying degrees. These observations suggest that neutralizing excess ROS may represent a broader strategy to mitigate disease manifestations associated with mitochondrial dysfunction and growth deficiency, including KS1. Future studies using our KS1 mouse model will be required to determine the extent to which antioxidant interventions can ameliorate the postnatal growth phenotype.

## Conclusions

Postnatal growth deficiency is one of the major hallmarks of KS1, but the mechanisms underlying this pathology are largely unknown. Longitudinal bone growth is driven by the proliferation, ECM production and hypertrophy of chondrocytes within the growth plate, ultimately leading to endochondral ossification and bone elongation. As chondrocytes mature, they undergo metabolic reprogramming from anaerobic glycolysis to OXPHOS to meet their increasing energy demands – a process that is supported by the progressively increasing environmental oxygen level. Here, we show that KMT2D-deficient chondrocytes, when exposed to supraphysiological oxygen levels, are unable to efficiently utilize mitochondrial respiration for energy production, leading to the accumulation of molecular oxygen, exacerbated ROS production, and a compensatory shift to anaerobic glycolysis as the major source for ATP generation. We found that the excessive ROS production accelerates the chondrocyte differentiation and maturation process, promoting their premature transition to the terminal hypertrophic, senescent state – a pathological process that can be successfully rescued with mitochondria-targeted antioxidants or the reduction of molecular oxygen, the source of mitochondrial ROS. Our study thus provides new evidence that oxygen-driven early growth plate senescence, which primarily affects the hypertrophic zone, is likely the underlying cause for the impairment in postnatal longitudinal bone growth in KS1.

## Methods

### Cell culture

ATDC5 cells were a gift from Prof. Jill A. Fahrner (Johns Hopkins University) (Fahrner et al., 2019). They were routinely cultured in DMEM/F12 medium supplemented with GlutaMAX (Gibco, 10565018), 5% FBS (Gibco, 10270106) and 1% penicillin/streptomycin in 5% CO2 in a 37°C cell culture incubator. Differentiation of ATDC5 cells was initiated as described before (Fahrner et al., 2019). Briefly, cells were seeded at ∼10,000 cells/cm^2^ and differentiation initiated 24 hours post-seeding (or 48 hours with MitoQ treatment) by adding the following supplements to the growth medium: 1x Insulin-Transferrin-Selenium (1x ITS; Thermo Fisher Scientific, 41400045), 50 µg/mL L-ascorbic acid (Sigma, A4544) and 10 mM Beta-glycerophosphate (Sigma, G9422). Medium was replaced every 2-3 days and differentiation carried out for up to 14 days. Hypoxia exposure was carried out in a Heracell VIOS 250i incubator (Thermo Fisher Scientific) in which the oxygen concentration was lowered by increasing the level of nitrogen (99.9%). Hypoxia exposure was administered at seeding and continued throughout differentiation.

### Histological staining

At various timepoints before and during the differentiation process, the cell cultures were fixed (4% paraformaldehyde) and stained with either 1% Alcian Blue in 3% acetic acid (Sigma-Aldrich, B8438) for 1 h, microscopically evaluated, staining extracted with 1% SDS for 20 min, and absorbance measured at 605 nm for quantification. On day 30 of differentiation, cultures were stained with 2% Alizarin Red (Abcam, ab146374) in ddH_2_O for 20 minutes. For quantification, Alizarin red was extracted with 10% Cetylpyridinium chloride for 10 min, the solution transferred to Eppendorf tubes, centrifuged at 16,100xg for 10 min, supernatant collected, and absorbance measured at 562 nm. Quantification was performed using SpectraMax^®^ M3 Microplate Reader (Molecular Devices) connected to LLX SoftMAx Pro 6.1 software.

### sc-RNA sequencing

sc-RNA-Seq was performed with Chromium Next GEM Single Cell 3’ Reagent Kits v3.1 (Dual Index) from 10XGenomics using RNA from ATDC5 cells. The experiment included four differentiation timepoints targeting 5.000 cells per sample following 10XGenomics protocols (Sample preparation demonstrated protocol – adherent cell lines and Chromium Next GEM Single Cell 3’ Reagent Kits v3.1, 2020, CG000315, Rev A). Briefly, cells were harvested at each timepoint, thoroughly resuspended in media, counted and filtered through a 100µm cell strainer (Falcon, CLS352235) to remove ECM followed by filtration through a 40 μm cell strainer (Pluriselect, 43-10040). Next, cells were centrifuged at 300xg for 3 minutes, resuspended in 1×PBS with 0.04% BSA, and counted. This step was repeated one more time and cells kept on ice while GEM ChipG were prepared according to the protocol. Cells were loaded at 1000 cells/µl according to the latest cell count, and cells captured at approx. 6000 cells per sample with Chromium Controller (10xGenomics). cDNA was kept at -20°C until the last timepoint and all libraries generated at the same time to eliminate additional technical variance. Libraries were generated with 25% of cDNA with recommended setup for 25-500 ng input of 12 cycle sample indexing. Both cDNA and library qualities were assessed with a Bioanalyzer 2100 High sensitivity chip (Agilent, 5067-4626). Libraries were pooled in two batches at 4 nM concentration and sequenced using NovaSeq6000 S1 at deCODE genetics.

### Hif1a knockdown

Cells were seeded as was previously described for differentiation and transfected with Lipofectamine 2000 (Thermo, 11668019) the following day. *Hif1a* knockdown was performed with two separate DsiRNA, siRNA1 (IDT, mm.Ri.Hif1a.13.1) or siRNA3 (IDT, mm.Ri.Hif1a.13.3) as well as a non-targeting control siRNA (IDT, 51-01-14-03). SiRNA:lipid complexes were re-introduced every other day with media replacement throughout differentiation, which was initiated 12 hours post-transfection. At each timepoint (days 0, 4, 7, and 14), Alcian Blue staining and RNA isolation were performed.

### Real time quantitative polymerase chain reaction (RT-qPCR)

RNA isolation was performed with a RNeasy Mini Kit (Qiagen, 74106) according to manufacturer’s protocol. Reverse transcription was performed with 0.4-1µg RNA input using High-Capacity cDNA Reverse Transcription Kit (Thermo, 4368814) and RT-qPCR performed using a Luna Universal qPCR Master Mix (NEB, M3003X) according to manufacturer’s protocols. The following primers were used: Hif1a primers (IDT, Mm.PT.58.11211292), Col10a1_F (5’-CATAAAGGGCCCACTTGCTA-3’), Col10a1_R (5’-CAGGAATGCCTTGTTCTCCT-3’), Hprt_F (5’-TCAGTCAACGGGGGACATAAA-3’), Hprt_R (5’-GGGGCTGTACTGCTTAACCAG-3’), Sox9_F (5’-CGGAACAGACTCACATCTCTCC-3’), Sox9_R (5’-GCTTGCACGTCGGTTTTGG-3’), Col2a1_F (5’-CGGTCCTACGGTGTCAGG-3’), Col2a1_R (5’-GCAGAGGACATTCCCAGTGT-3’). RT-qPCR was performed on a CFX384 Real-Time System (Bio-Rad), and analysis performed with CFXmanager software (Bio-Rad, v. 3.1).

### Metabolic flux measurements

For the assessment of cellular respiration, XF Cell Mito Stress test was performed using a Seahorse XFe96 FluxPaks (Seahorse Bioscience, 102416-100) according to manufacturer’s protocol. Briefly, 5.000 cells/well were seeded the day before the experiment in normal growth medium and sensors hydrated in calibration buffer at 37°C. On the day of the experiment, cells were washed with 1×PBS and running media added composed of Seahorse XF DMEM medium, pH 7.4 (Agilent, 103575-100), 1 mM pyruvate solution (Agilent, 103578-100) and 10 mM D-(+)-glucose (Sigma, G7021) for 1 hour in 37°C without CO_2_. In addition, for the galactose experiment, the running media was supplemented with 10 mM galactose (Sigma, G5388) instead of glucose and cells incubated for 1 hour with or without galactose. Respiration was measured with XF96 Extracellular Flux Analyzer (Seahorse Bioscience, Billerica, MA, USA) according to the manufacturer’s instructions for basal, maximal and spare respiration. Basal oxygen consumption rate (OCR) was measured with six consecutive measurements at 6.5 min intervals. This was followed by three measurements after the addition of each mitochondrial inhibitor, oligomycin (1.5μM) and carbonyl cyanide p-triflouromethoxyphenylhydrazone (FCCP) (1.5μM) to induce maximal respiration. Finally, a combination of rotenone/antimycin A (both at 1.25μM) was injected and three consecutive measurements were taken for the calculation of spare respiration. Normalization was performed with either Presto blue or Hoechst. Before starting the Mito Stress test, Presto blue (Invitrogen, A13262) was added and incubated for 1 hour. Media with the dye was then moved to a new plate, and cells washed with 1×PBS before running the Mito Stress test. Absorbance of Presto blue was measured at 570 and 600 nm using a microplate reader (SpectraMax^®^ M3, Molecular Devices) and used for cell count/seeding density corrections. In the galactose experiment, 2μM Hoechst 33342 (Thermo, 62249) was added after the assay for 30 min, cells were harvested and moved to a black 96-well plate, and the fluorescence was measured with GloMax (Promega).

Lactate measurements were performed using spent media with ABL800FLEX (Radiometer, Brønshøj, Denmark). Media was collected from both *Kmt2d^+/^* and *Kmt2d^-/^* cells on days 0, 4, 7 or 14 of differentiation with addition of a well containing no cells as baseline measurement control. Media was collected at individual timepoints, frozen at -80°C and all samples measured at the same time.

### Fluorescent dye-based assays for the measurement of cell division, ROS production, and cellular senescence

A CellTrace™ CFSE Cell Proliferation Kit (Thermo, C34554) was used to assess proliferation. Cells were harvested and 750.000 cells resuspended in 1 ml 1×PBS. Next, CFSE was added at a 10 µM final concentration and incubated for 20 minutes. The staining was diluted with 5 ml growth media and incubated for an additional 5 minutes. Cells were centrifuged at 300x g for 5 minutes, resuspended in 1 ml growth medium, counted, and 125.000 cells were seeded into a T25 cell culture dish for expansion. Leftover cells were used to measure baseline fluorescence. After 72 hours of expansion, cells were collected by trypsinization, washed, centrifuged at 300x g for 5 minutes, and resuspended in FACS buffer (1×PBS with 2% FBS) before passing through a cell strainer. Fluorescence was measured using a flow cytometer (Attune NxT Flow Cytometer).

Superoxide (•O_2_^-^) levels were measured on day 0 using MitoSox (Stemcell, cat. 100-0991). Cells were stained with MitoSox (1:1000) in HBSS (Thermo, 14025092) for 30 minutes at 37°C, 5% CO_2_. After incubation, staining solution was removed and cells washed with HBSS. Next, cells were harvested by trypsinization, trypsin diluted with FACS buffer (1×PBS with 2% FBS), cells passed through a cell strainer and fluorescence measured using a flow cytometer (Attune NxT Flow Cytometer).

Cellular hydrogen peroxide (H_2_O_2_) was measured on days 0, 4 and 7 with 2′,7′-Dichlorofluorescin Diacetate (DCFDA) (Sigma, 287810). On the day of measurement, the media was removed, 20 µM DCFDA in appropriate media (maintenance or differentiation) added to the well and incubated for 30 minutes at 37°C, 5% CO_2_ in the presence of either 20% or 5% O_2_. At the end of incubation, the staining solution was removed, and the cells were washed with 1×PBS, harvested by trypsinization, diluted with FACS buffer (1×PBS with 2% FBS), filtered through a cell strainer before their fluorescence was measured using a flow cytometer (Attune NxT Flow Cytometer).

β-Galactosidase activity was measured using a Senescence Assay Kit (Abcam, ab228562). On the day of the assay, i.e., on differentiation day 20, fresh medium was added containing 1.5 µl senescence dye per 500 µl medium, and incubated at 37°C, 5% CO_2_ for 2 hours. After the incubation, the cells were washed twice with Assay Buffer XXVII/Wash Buffer, and the cells were harvested by trypsinization. Cells were centrifuged at 300xg for 5 minutes, resuspended in Assay Buffer XXVII/Wash Buffer, and the fluorescence was measured using a flow cytometer (Attune NxT Flow Cytometer).

### Mice

C57BL/6J *Kmt2d^+/βGeo^* mice have been previously described (Fahrner et al., 2019). Mice were housed in the clean, specific pathogen-free state-of-the-art ArcticLAS animal facility (Reykjavik, Iceland) in ventilated racks and provided ad libitum access to a standard rodent diet and to filtered water. All female C57BL/6J wild-type mice used in this study were between 6 weeks and 3 months old. Import and all experimental protocols were approved by the Icelandic Food and Veterinary Authority (license: 18081393) and approved by the National Expert Advisory Board on Animal Welfare.

### *In vivo* staining

C57BL/6J wild-type female mice were mated with C57BL/6J *Kmt2d^+/βGeo^* males overnight. Pregnant dams were weighed and the nitroimidazole hypoxia marker EF5 (Medchemexpress, HY-U00118) was administered through intraperitoneal injection with 10 mM EF5 at 1% of body weight (0.1 mL per 10 g weight) 17.5 days post-conception. Three hours later, the E17.5 embryos were dissected in cold 1×PBS, rinsed in 1×PBS, and the hindlimbs were fixed for 2 hours with 4% PFA, incubated overnight in 30% sucrose, and embedded in OCT. Tissue was sectioned (10 µm) on a Leica CM1850 cryostat using Kawamoto’s adhesive film method to maintain tissue integrity (Kawamoto T., 2003). Sections were stored at −80°C until staining. The first to third sections into the bone marrow were used for the staining.

Sections were permeabilized for 10 minutes at 4°C with acetone, adding acetone every 3 minutes to counteract evaporation. The sections were then aspirated and rinsed once with TBS. Sections were incubated for 1 hour at room temperature with blocking buffer (TBS with 0.05% triton-X 100 and 5% normal goat serum). Primary antibodies (1:200 unconjugated Anti-Collagen X, Abcam #ab260040; 1:100 Anti-EF5 Cy5 conjugate, Merck Millipore, #EF5012) were diluted in blocking buffer, then added to the sections and incubated overnight at 4°C in humidity chamber. Sections were washed with wash buffer (TBS with 0.05% triton-X 100) three times for 10 minutes at room temperature. Secondary antibody (Goat anti-Mouse IgG (H+L) Secondary Antibody Alexa Fluor 555, Thermo #A32727) to detect Collagen X was diluted 1:1,000 in blocking buffer, before being added to sections and incubated overnight at 4°C in humidity chamber. Slides were washed with wash buffer three times for 10 minutes, and once with TBS for 10 minutes at room temperature, before being mounted with Fluoromount G with DAPI (Thermo #00-4959-52). Slides were stored for 24 hours at 4°C, exposed to air, and then stored at 4°C prior to imaging.

Slides were imaged with an Olympus FV4000 confocal using 10x objective, using a step size of 1.5µm. Stitching was performed using the cellSens FluoView software. Fiji was used to generate maximum intensity projections and analysis (Schindelin et al., 2012). The hypertrophic and hypoxic regions of the distal femur growth plate, as well as the total growth plate (from the top of the resting zone to the edge of the hypertrophic zone), were manually traced using Fiji software. The student performing the sectioning, staining, and analysis was blinded to the genotype. Each data point represents a biological replicate.

## Data analysis

Statistical analysis was performed in GraphPad Prism (v.10.1.1) and RNA sequencing analysis in Rstudio. Alignment and matrix generation for sc-RNA sequencing was done with CellRanger (v. 6.1.2; (G. X. Y. Zheng et al., 2017)) using default settings and the mm10 reference genome provided by 10x Genomics (2020). Output h5 files were imported to R studio (v. 4.3.0) with Seurat (v. 5.1.0; (Hao et al., 2024)) and each sample filtered according to nFeature_RNA > 200 and percent.mt < 20. Samples were individually normalized using Seurat’s NormalizeData function with default settings before merging. Highly variable features were identified using Seurat’s FindVariableFeatures, and genes scaled with ScaleData. Principal component analysis (PCA) was then performed with 50 components and Harmony used for batch correction (v. 1.2.0; (Korsunsky et al., 2019)). Uniform manifold approximation and projection (UMAP) was run on the Harmony-reduced dimensions (1:20) and cell clusters calculated with FindNeighbors and FindClusters (0.5 resolution). Identity markers for each cluster were found with FindAllMarkers. For cell cycle analysis, the Seurat object was converted using SingleCellExperiment (v. 1.24.0; (Amezquita et al., 2020)) and phase predictions performed with Tricycle (v.1.10.0; (S. C. Zheng et al., 2022)) using project_cycle_space and estimate_cycle_position. Slingshot (v. 2.10.0; (Street et al., 2018)) was used for differentiation trajectory inference with UMAP embeddings for cellular transition model.

Previously published bulk RNA sequencing Fastq files from ATDC5 (Fahrner et al., 2019) were accessed through the EMBL-EBI ENA database (PRJNA531019) and pseudo-aligned to the GRCm39 mouse transcriptome, downloaded from Ensembl, using Kallisto (v.0.48.0; (Bray et al., 2016)). Output files were imported to R studio (v. 4.3.0) using Tximport (v. 1.30.0; (Soneson et al., 2015)). Transcripts were assigned with TxDb.Mmusculus.UCSC.mm10.ensGene (v.3.4.0). Before analysis, transcripts with ≤10 total counts across all samples and genes expressed in ≤6 samples were filtered out. Differential expression analysis was performed with DESeq2 (v.1.42.1; (Love et al., 2014)) and GSEA done with fgsea package (v. 1.28.0; (Korotkevich et al., 2016)) using only differentially expressed genes with a baseMean > 10 and log2 fold change above 1 or below -1. GSEA was performed using minSize=15, maxSize=1000 and Hallmark gene set (MH) downloaded from the Mouse Molecular Signatures Database (MSigDB).

## Author contributions

S.T.H., H.T.B., and A.U. designed and directed the study. H.T.B. received the funding for the study. S.T.H. and A.U. wrote the initial manuscript draft. S.T.H. and S.P. performed experiments and acquired the data. S.T.H., H.T.B., A.U., and S.P. analyzed the data. All authors edited and read the final version of the manuscript.

## Acknowledgments

We thank Dr. Romain Lasseur for the help with mouse breeding and EF5 injections, the ArcticLAS animal facility for technical assistance, and Amgen deCODE genetics for helping with a subset of the sequencing.

## Figures

**Supplementary Figure 1:**
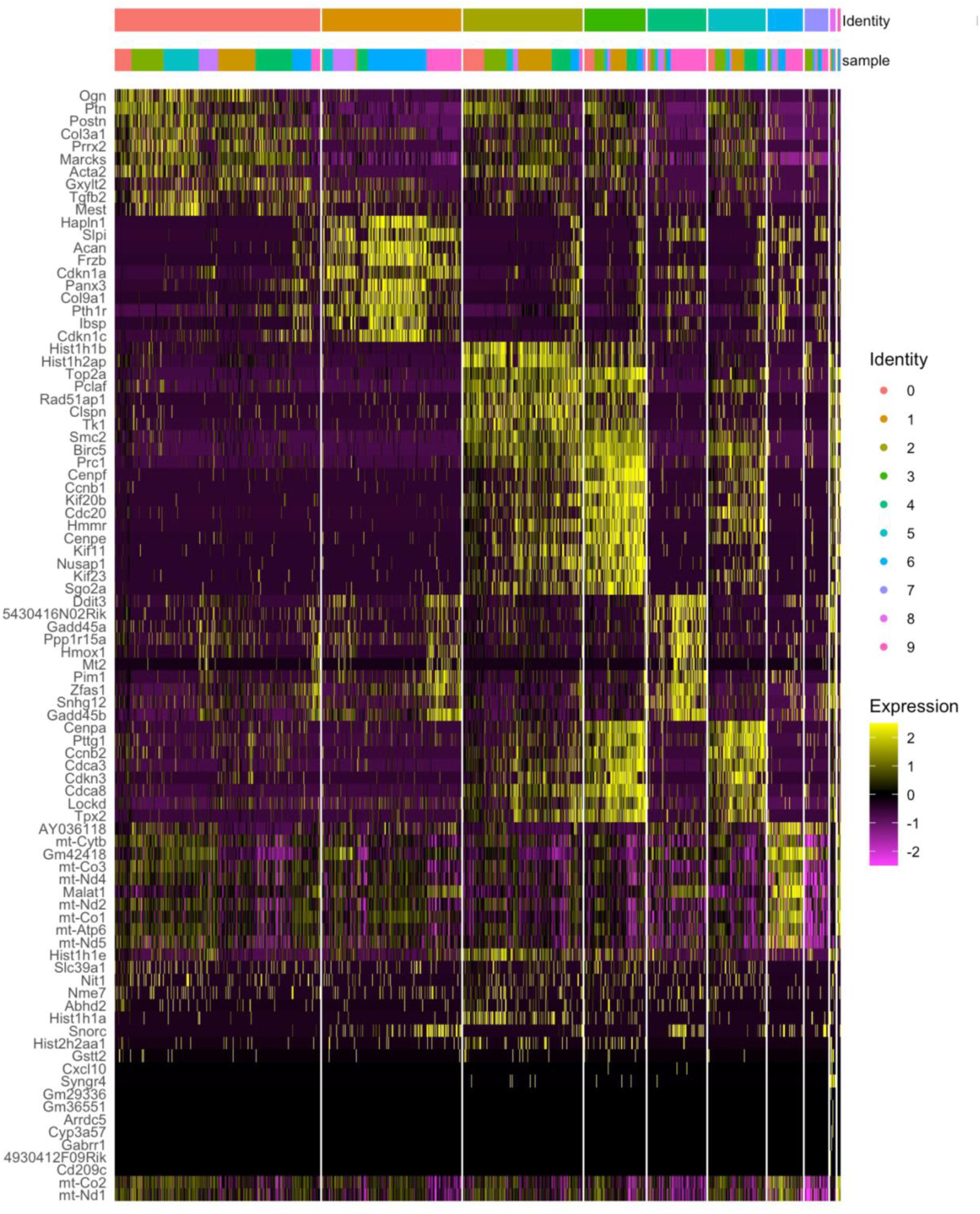
Heatmap showing the top 10 positively enriched marker genes for each of the 10 clusters identified by single-cell RNA sequencing. Columns represent individual cells, grouped and annotated by cluster identity, and the rows correspond to marker genes ranked by expression within each cluster. The top annotation bars indicate cluster assignment and sample origin. Color intensity reflects gene expression, with higher expression shown in yellow and lower expression in purple.

**Supplementary Figure 2:**
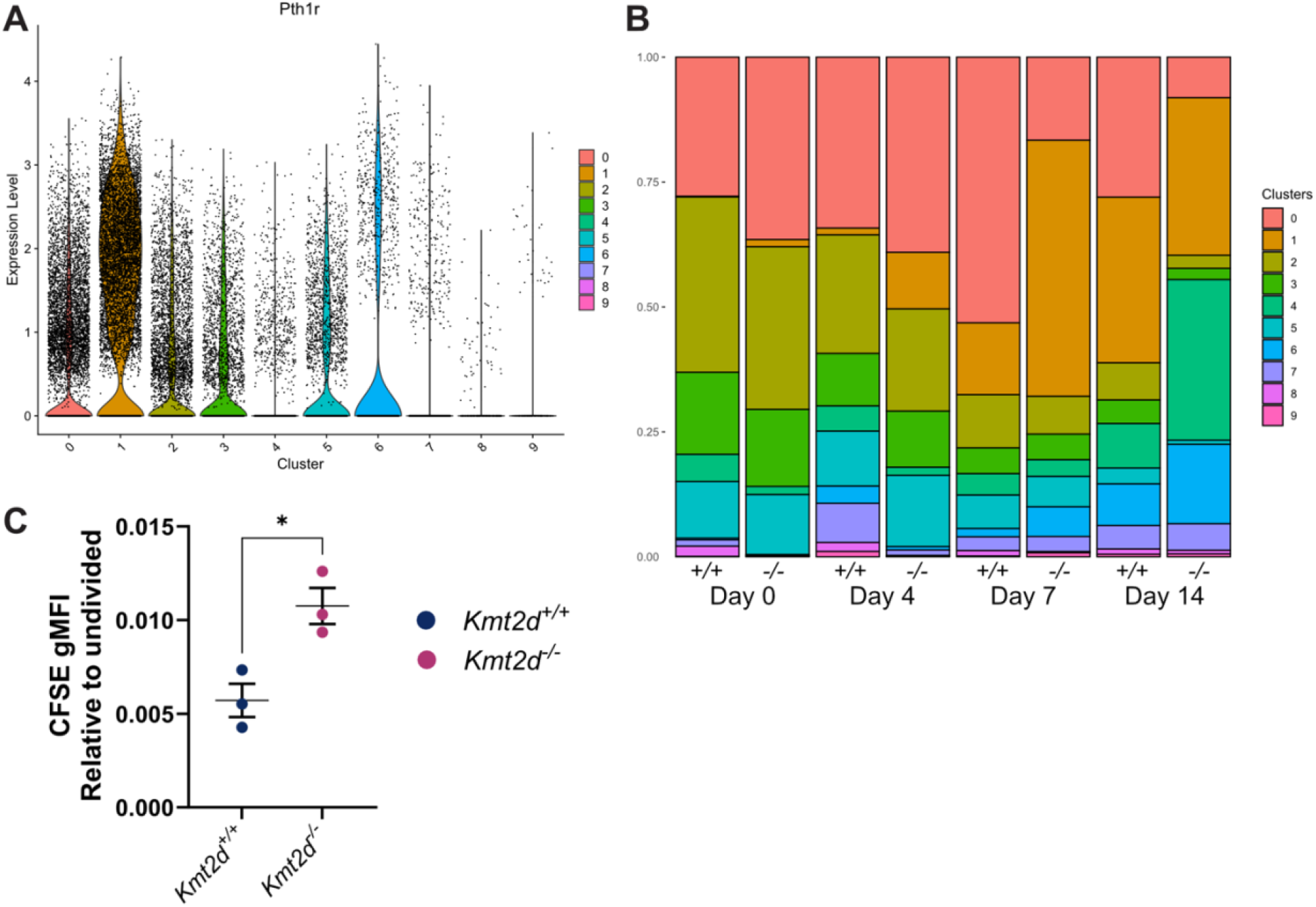
A) Violin plot of *Pth1r* expression in sc-RNA sequencing data across clusters. B) sc-RNA sequencing cell count distributions across clusters within each sample. C) CFSE fluorescence for cell division measured 72 hours post-seeding and normalized to fluorescence of the corresponding undivided cells. Data shown as mean ± SEM and statistical test performed with unpaired Student’s t-test, *P ≤ 0.05.

**Supplementary Figure 3:**
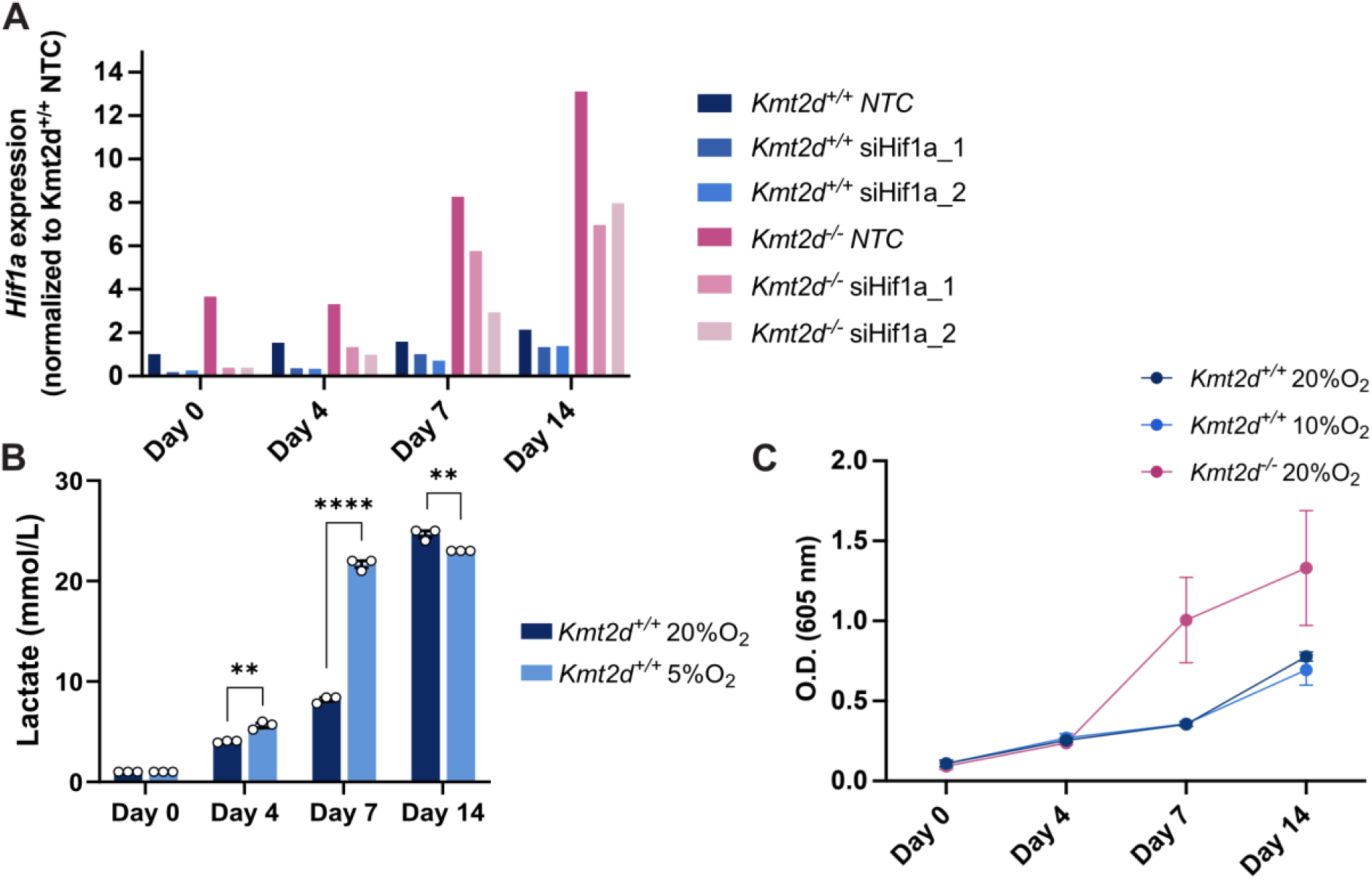
A) *Hif1a* expression in *Kmt2d^+/+^* and *Kmt2d^-/-^* cells upon treatment with the *Hif1a*-targeting siRNAs siHif1a_1 and siHif1a_2, or non-targeting control (NTC) siRNA, normalized to K*mt2d^+/+^* NTC cells throughout 14-day differentiation. B) Amperometry measurements for lactate in mmol/L from culture media of *Kmt2d^+/+^* cells at four timepoints during differentiation in the presence of either 20% or 5% environmental O_2_. Data shown as mean ± SEM and statistical tests performed with unpaired Student’s t-tests within each timepoint. **P ≤ 0.01, ****P ≤ 0.0001. C) Quantification of Alcian blue staining for *Kmt2d^-/-^* cells differentiated in the presence of 20% environmental O_2_ and *Kmt2d^+/+^* cells differentiated in the presence of either 20% or 10% environmental O_2._

**Supplementary Figure 4:**
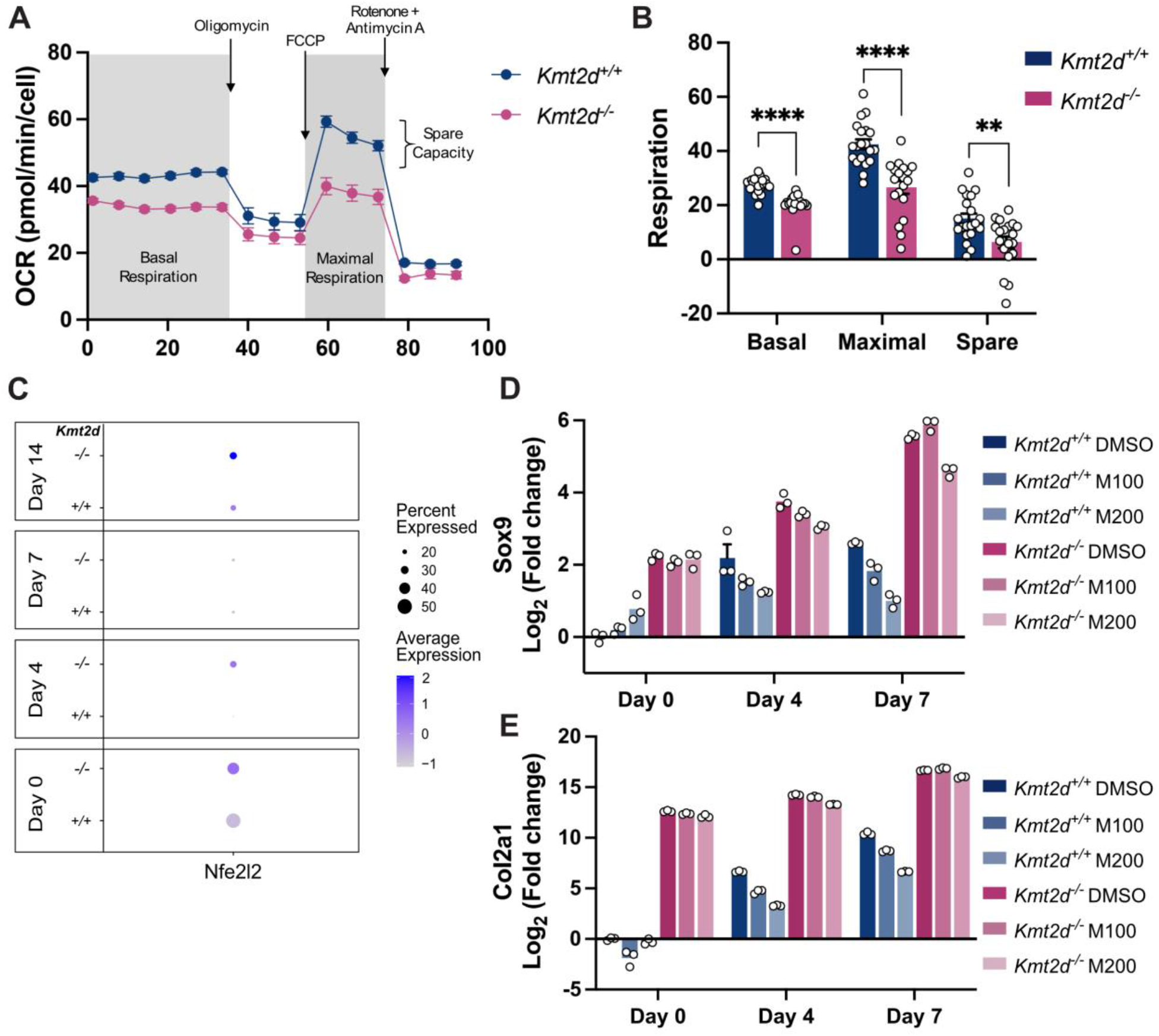
Oxygen consumption rate (OCR) of the second biological replicate of undifferentiated *Kmt2d^+/+^* and *Kmt2d^-/-^* cells measured with Seahorse assay. A) Average respiration rate across all technical replicates over time. B) Mitochondria-specific basal and maximal respiration were calculated by subtracting the non-mitochondrial respiration from the total basal and maximal respiration, respectively. Spare respiration was calculated by subtracting the mitochondria-specific basal respiration from the mitochondria-specific maximal respiration. Data shown as mean ± SEM. Statistical tests were performed with unpaired Student’s t-tests. **P ≤ 0.01, ****P ≤ 0.0001. C) A DotPlot generated from the sc-RNA-Seq data displaying the percentage of *Kmt2d^+/+^* and *Kmt2d^-/-^* cells expressing *Nfe2l2* and the average *Nfe2l2* expression across all cells at four different timepoints during the differentiation process. D) *Sox9* and E) *Col2a1* expression of *Kmt2d^+/+^* and *Kmt2d^-/-^* cells with or without MitoQ treatment during differentiation.

**Supplementary Figure 5:**
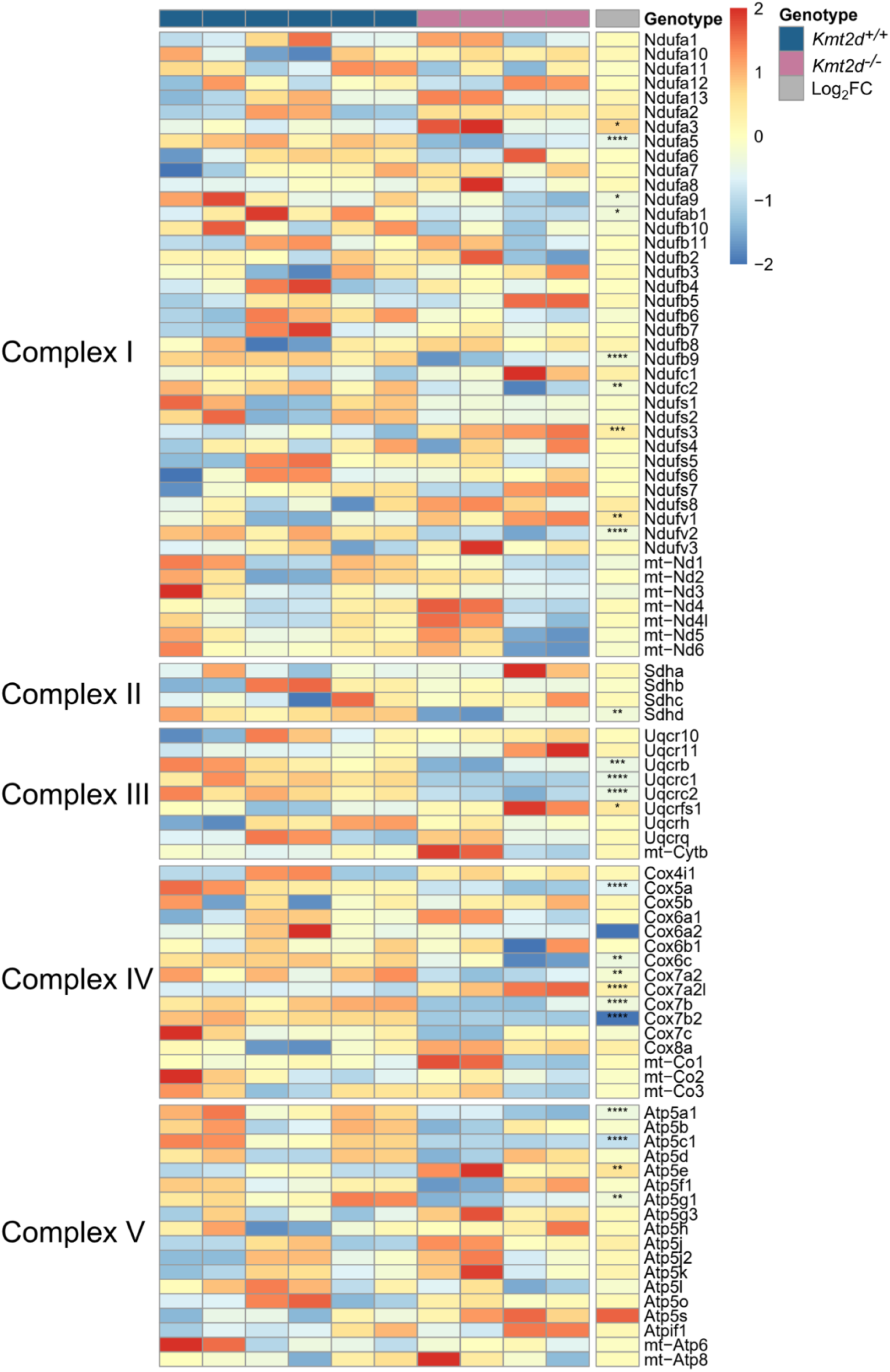
Heatmap of the expression levels of 90 mitochondrial and nuclear encoded genes for electron transport chain (ETC) complex subunits in undifferentiated chondrocytes. Genes are organized by ETC complex (I–V), and samples are grouped by genotype, with *Kmt2d^+/+^* labeled in blue and *Kmt2d^-/-^* in pink. Color intensity represents row-scaled expression values across samples, with higher expression shown in red and lower expression in blue. The final column labeled in grey, indicates the log_2_ fold change, with increased expression in *Kmt2d^-/-^* relative to *Kmt2d^+/+^* shown in red and decreased expression in blue. Significance of differential expression is indicated by asterisks: adjusted P value(padj) * ≤ 0.05, ** ≤ 0.01, *** ≤ 0.001, **** ≤ 0.0001.

## References

Adam, M. P., Banka, S., Bjornsson, H. T., Bodamer, O., Chudley, A. E., Harris, J., Kawame, H., Lanpher, B. C., Lindsley, A. W., Merla, G., Miyake, N., Okamoto, N., Stumpel, C. T., & Niikawa, N. (2019). Kabuki syndrome: international consensus diagnostic criteria. Journal of Medical Genetics, 56(2), 89–95. 10.1136/jmedgenet-2018-105625

Amezquita, R. A., Lun, A. T. L., Becht, E., Carey, V. J., Carpp, L. N., Geistlinger, L., Marini, F., Rue-Albrecht, K., Risso, D., Soneson, C., Waldron, L., Pagès, H., Smith, M. L., Huber, W., Morgan, M., Gottardo, R., & Hicks, S. C. (2020). Orchestrating single-cell analysis with Bioconductor. Nature Methods, 17(2), 137–145. 10.1038/s41592-019-0654-x

Araldi, E., & Schipani, E. (2010). Hypoxia, HIFs and bone development. Bone, 47(2), 190–196. 10.1016/j.bone.2010.04.606

Arnold, M. A., Kim, Y., Czubryt, M. P., Phan, D., McAnally, J., Qi, X., Shelton, J. M., Richardson, J. A., Bassel-Duby, R., & Olson, E. N. (2007). MEF2C Transcription Factor Controls Chondrocyte Hypertrophy and Bone Development. Developmental Cell, 12(3), 377–389. 10.1016/j.devcel.2007.02.004

Bakare, A. B., Lesnefsky, E. J., & Iyer, S. (2021). Leigh Syndrome: A Tale of Two Genomes. In Frontiers in Physiology (Vol. 12). Frontiers Media S.A. 10.3389/fphys.2021.693734

Blumer, M. J. F. (2021). Bone tissue and histological and molecular events during development of the long bones. Annals of Anatomy, 235, 151704. 10.1016/j.aanat.2021.151704

Bolduc, J. A., Collins, J. A., & Loeser, R. F. (2019). Reactive oxygen species, aging and articular cartilage homeostasis. Free Radical Biology and Medicine, 132(April 2018), 73–82. 10.1016/j.freeradbiomed.2018.08.038

Bray, N. L., Pimentel, H., Melsted, P., & Pachter, L. (2016). Near-optimal probabilistic RNA-seq quantification. Nature Biotechnology, 34(5), 525–527. 10.1038/nbt.3519

Carosso, G. A., Boukas, L., Augustin, J. J., Nguyen, H. N., Winer, B. L., Cannon, G. H., Robertson, J. D., Zhang, L., Hansen, K. D., Goff, L. A., & Bjornsson, H. T. (2019). Precocious neuronal differentiation and disrupted oxygen responses in Kabuki syndrome. JCI Insight, 4(20), 1–17. 10.1172/jci.insight.129375

Chandel, N. S. (2014). Mitochondria as signaling organelles. In BMC Biology (Vol. 12). BioMed Central Ltd. 10.1186/1741-7007-12-34

Chatzi, C., Schnell, E., & Westbrook, G. L. (2015). Localized hypoxia within the subgranular zone determines the early survival of newborn hippocampal granule cells. ELife, 4(OCTOBER2015), 1–18. 10.7554/eLife.08722

Cho, J. H., Jung, H. W., & Shim, K. S. (2024). Growth plate closure and therapeutic interventions. Clinical and Experimental Pediatrics, 67(11), 553–559. 10.3345/cep.2023.00346

Chu, T. L., Chen, P., Yu, A. X., Kong, M., Tan, Z., Tsang, K. Y., Zhou, Z., & Cheah, K. S. E. (2023). MMP14 cleaves PTH1R in the chondrocyte-derived osteoblast lineage, curbing signaling intensity for proper bone anabolism. ELife, 12, 1–28. 10.7554/elife.82142

Cline, S. D. (2012). Mitochondrial DNA damage and its consequences for mitochondrial gene expression. In Biochimica et Biophysica Acta - Gene Regulatory Mechanisms (Vol. 1819, Numbers 9–10, pp. 979–991). 10.1016/j.bbagrm.2012.06.002

Cvetko, F., Caldwell, S. T., Higgins, M., Suzuki, T., Yamamoto, M., Prag, H. A., Hartley, R. C., Dinkova-Kostova, A. T., & Murphy, M. P. (2021). Nrf2 is activated by disruption of mitochondrial thiol homeostasis but not by enhanced mitochondrial superoxide production. Journal of Biological Chemistry, 296(24), 100169. 10.1074/jbc.RA120.016551

Eruslanov, E., & Kusmartsev, S. (2010). Identification of ROS Using Oxidized DCFDA and Flow-Cytometry BT - Advanced Protocols in Oxidative Stress II (D. Armstrong, Ed.; pp. 57–72). Humana Press. 10.1007/978-1-60761-411-1_4

Fahrner, J. A., Lin, W. Y., Riddle, R. C., Boukas, L., DeLeon, V. B., Chopra, S., Lad, S. E., Luperchio, T. R., Hansen, K. D., & Bjornsson, H. T. (2019). Precocious chondrocyte differentiation disrupts skeletal growth in Kabuki syndrome mice. JCI Insight, 4(20), 1–19. 10.1172/jci.insight.129380

Froimchuk, E., Jang, Y., & Ge, K. (2017). Histone H3 lysine 4 methyltransferase KMT2D. Gene, 627(June), 337–342. 10.1016/j.gene.2017.06.056

Hamanaka, R. B., & Chandel, N. S. (2010). Mitochondrial reactive oxygen species regulate cellular signaling and dictate biological outcomes. In Trends in Biochemical Sciences (Vol. 35, Number 9, pp. 505–513). 10.1016/j.tibs.2010.04.002

Hao, Y., Stuart, T., Kowalski, M. H., Choudhary, S., Hoffman, P., Hartman, A., Srivastava, A., Molla, G., Madad, S., Fernandez-Granda, C., & Satija, R. (2024). Dictionary learning for integrative, multimodal and scalable single-cell analysis. Nature Biotechnology, 42(2), 293–304. 10.1038/s41587-023-01767-y

Hollander, J. M., Li, L., Rawal, M., Wang, S. K., Shu, Y., Zhang, M., Nielsen, H. C., Rosen, C. J., & Zeng, L. (2022). A critical bioenergetic switch is regulated by IGF2 during murine cartilage development. Communications Biology, 5(1), 1–13. 10.1038/s42003-022-04156-4

Holzer, T., Probst, K., Etich, J., Auler, M., Georgieva, V. S., Bluhm, B., Frie, C., Heilig, J., Niehoff, A., Nüchel, J., Plomann, M., Seeger, J. M., Kashkar, H., Baris, O. R., Wiesner, R. J., & Brachvogel, B. (2019). Respiratory chain inactivation links cartilage-mediated growth retardation to mitochondrial diseases. Journal of Cell Biology, 218(6), 1853–1870. 10.1083/JCB.201809056

Kauffman, M., Kauffman, M., Traore, K., Zhu, H., Trush, M., Jia, Z., & Li, Y. (2016). MitoSOX-Based Flow Cytometry for Detecting Mitochondrial ROS. Reactive Oxygen Species, 2(5), 361–370. 10.20455/ros.2016.865

Kawamoto T. (2003). Use of a new adhesive film for the preparation of multi-purpose fresh-frozen sections from hard tissues, whole-animals, insects and plants. Archives of Histology and Cytology, 66(2), 123–143. 10.1679/aohc.66.123

Ki, S. K., Hae, W. C., Hee, E. Y., & Ick, Y. K. (2010). Reactive oxygen species generated by NADPH oxidase 2 and 4 are required for chondrogenic differentiation. Journal of Biological Chemistry, 285(51), 40294–40302. 10.1074/jbc.M110.126821

Korotkevich, G., Sukhov, V., Budin, N., Shpak, B., Artyomov, M. N., & Sergushichev, A. (2016). Fast gene set enrichment analysis. BioRxiv. 10.1101/060012

Korsunsky, I., Millard, N., Fan, J., Slowikowski, K., Zhang, F., Wei, K., Baglaenko, Y., Brenner, M., Loh, P. ru, & Raychaudhuri, S. (2019). Fast, sensitive and accurate integration of single-cell data with Harmony. Nature Methods, 16(12), 1289–1296. 10.1038/s41592-019-0619-0

Long, F., & Ornitz, D. M. (2013). Development of the endochondral skeleton. Cold Spring Harbor Perspectives in Biology, 5(1). 10.1101/cshperspect.a008334

Love, M. I., Huber, W., & Anders, S. (2014). Moderated estimation of fold change and dispersion for RNA-seq data with DESeq2. Genome Biology, 15(12). 10.1186/s13059-014-0550-8

Lv, S., Wen, H., Shan, X., Li, J., Wu, Y., Yu, X., Huang, W., & Wei, Q. (2019). Loss of KMT2D induces prostate cancer ROS-mediated DNA damage by suppressing the enhancer activity and DNA binding of antioxidant transcription factor FOXO3. Epigenetics, 14(12), 1194–1208. 10.1080/15592294.2019.1634985

Mao, H., Zhang, Y., Xiong, Y., Zhu, Z., Wang, L., & Liu, X. (2022). Mitochondria-Targeted Antioxidant Mitoquinone Maintains Mitochondrial Homeostasis through the Sirt3-Dependent Pathway to Mitigate Oxidative Damage Caused by Renal Ischemia/Reperfusion. Oxidative Medicine and Cellular Longevity, 2022. 10.1155/2022/2213503

Monti, L., Paganini, C., Lecci, S., De Leonardis, F., Hay, E., Cohen-Solal, M., Villani, S., Superti-Furga, A., Tenni, R., Forlino, A., & Rossi, A. (2015). N-acetylcysteine treatment ameliorates the skeletal phenotype of a mouse model of diastrophic dysplasia. Human Molecular Genetics, 24(19), 5570–5580. 10.1093/hmg/ddv289

Morita, K., Miyamoto, T., Fujita, N., Kubota, Y., Ito, K., Takubo, K., Miyamoto, K., Ninomiya, K., Suzuki, T., Iwasaki, R., Yagi, M., Takaishi, H., Toyama, Y., & Suda, T. (2007). Reactive oxygen species induce chondrocyte hypertrophy in endochondral ossification. Journal of Experimental Medicine, 204(7), 1613–1623. 10.1084/jem.20062525

Murphy, M. P. (2001). Development of lipophilic cations as therapies for disorders due to mitochondrial dysfunction. Expert Opinion on Biological Therapy, 1(5), 753–764. 10.1517/14712598.1.5.753

Newton, P. T., Staines, K. A., Spevak, L., Boskey, A. L., Teixeira, C. C., Macrae, V. E., Canfield, A. E., & Farquharson, C. (2012). Chondrogenic ATDC5 cells: An optimised model for rapid and physiological matrix mineralisation. International Journal of Molecular Medicine, 30(5), 1187–1193. 10.3892/ijmm.2012.1114

Pacelli, C., Adipietro, I., Malerba, N., Squeo, G. M., Piccoli, C., Amoresano, A., Pinto, G., Pucci, P., Lee, J. E., Ge, K., Capitanio, N., & Merla, G. (2020). Loss of Function of the Gene Encoding the Histone Methyltransferase KMT2D Leads to Deregulation of Mitochondrial Respiration. Cells, 9(7), 1–20. 10.3390/cells9071685

Posey, K. L., Coustry, F., Veerisetty, A. C., Hossain, M., Alcorn, J. L., & Hecht, J. T. (2015). Antioxidant and anti-inflammatory agents mitigate pathology in a mouse model of pseudoachondroplasia. Human Molecular Genetics, 24(14), 3918–3928. 10.1093/hmg/ddv122

Puchtler, H., Meloan, S. N., & Terry, M. S. (1969). On the history and mechanism of alizarin and alizarin red S stains for calcium. Journal of Histochemistry & Cytochemistry., 17(2), 110–124. 10.1177/17.2.110

Schindelin, J., Arganda-Carreras, I., Frise, E., Kaynig, V., Longair, M., Pietzsch, T., Preibisch, S., Rueden, C., Saalfeld, S., Schmid, B., Tinevez, J. Y., White, D. J., Hartenstein, V., Eliceiri, K., Tomancak, P., & Cardona, A. (2012). Fiji: An open-source platform for biological-image analysis. In Nature Methods (Vol. 9, Number 7, pp. 676–682). 10.1038/nmeth.2019

Schipani, E., Ryan, H. E., Didrickson, S., Kobayashi, T., Knight, M., & Johnson, R. S. (2001). Hypoxia in cartilage: HIF-1α is essential for chondrocyte growth arrest and survival. Genes and Development, 15(21), 2865–2876. 10.1101/gad.934301

Schott, D. A., Blok, M. J., Gerver, W. J. M., Devriendt, K., Zimmermann, L. J. I., & Stumpel, C. T. R. M. (2016). Growth pattern in Kabuki syndrome with a KMT2D mutation. American Journal of Medical Genetics, Part A, 170(12), 3172–3179. 10.1002/ajmg.a.37930

Soneson, C., Love, M. I., & Robinson, M. D. (2015). Differential analyses for RNA-seq: transcript-level estimates improve gene-level inferences. F1000Research, *4*, 1521. 10.12688/f1000research.7563.1

Stegen, S., Laperre, K., Eelen, G., Rinaldi, G., Fraisl, P., Torrekens, S., Van Looveren, R., Loopmans, S., Bultynck, G., Vinckier, S., Meersman, F., Maxwell, P. H., Rai, J., Weis, M., Eyre, D. R., Ghesquière, B., Fendt, S. M., Carmeliet, P., & Carmeliet, G. (2019). HIF-1α metabolically controls: collagen synthesis and modification in chondrocytes. Nature, 565(7740), 511–515. 10.1038/s41586-019-0874-3

Street, K., Risso, D., Fletcher, R. B., Das, D., Ngai, J., Yosef, N., Purdom, E., & Dudoit, S. (2018). Slingshot: Cell lineage and pseudotime inference for single-cell transcriptomics. BMC Genomics, 19(1). 10.1186/s12864-018-4772-0

Ulfig, A., & Jakob, U. (2024). Cellular oxidants and the proteostasis network: balance between activation and destruction. Trends in Biochemical Sciences, 49(9), 761–774. 10.1016/j.tibs.2024.07.001

Vega, R. B., Matsuda, K., Oh, J., Barbosa, A. C., Yang, X., Meadows, E., McAnally, J., Pomajzl, C., Shelton, J. M., Richardson, J. A., Karsenty, G., & Olson, E. N. (2004). Histone deacetylase 4 controls chondrocyte hypertrophy during skeletogenesis. Cell, 119(4), 555–566. 10.1016/j.cell.2004.10.024

Yao, Q., Khan, M. P., Merceron, C., LaGory, E. L., Tata, Z., Mangiavini, L., Hu, J., Vemulapalli, K., Chandel, N. S., Giaccia, A. J., & Schipani, E. (2019). Suppressing Mitochondrial Respiration Is Critical for Hypoxia Tolerance in the Fetal Growth Plate. Developmental Cell, 49(5), 748–763.e7. 10.1016/j.devcel.2019.04.029

Zheng, G. X. Y., Terry, J. M., Belgrader, P., Ryvkin, P., Bent, Z. W., Wilson, R., Ziraldo, S. B., Wheeler, T. D., McDermott, G. P., Zhu, J., Gregory, M. T., Shuga, J., Montesclaros, L., Underwood, J. G., Masquelier, D. A., Nishimura, S. Y., Schnall-Levin, M., Wyatt, P. W., Hindson, C. M., … Bielas, J. H. (2017). Massively parallel digital transcriptional profiling of single cells. Nature Communications, 8. 10.1038/ncomms14049

Zheng, S. C., Stein-O’Brien, G., Augustin, J. J., Slosberg, J., Carosso, G. A., Winer, B., Shin, G., Bjornsson, H. T., Goff, L. A., & Hansen, K. D. (2022). Universal prediction of cell-cycle position using transfer learning. Genome Biology, 23(1). 10.1186/s13059-021-02581-y

